# Pre-existing intratumoral stem-like CD8^+^ T-cells drive radiotherapy-induced tumor control

**DOI:** 10.1101/2024.10.16.618635

**Authors:** Hakan Köksal, Michael Herbst, Paulo Perreira, Marc Nater, Nicola Regli, Cheïma Boudjeniba, Nese Erdem Borgoni, Virginia Cecconi, Maries van den Broek

## Abstract

CD8^+^ T-cells are essential for spontaneous and therapy-induced restriction of tumor progression. Upon radiotherapy, these pre-existing CD8^+^ T-cells expand rapidly after their initial decline and are essential and sufficient for early tumor control. Stem-like CD8^+^ T-cells proliferate and differentiate into effector CD8^+^ T-cells in response to radiotherapy and are thus key determinants of therapy efficacy. Our findings highlight the critical role of intratumoral stem-like CD8^+^ T-cells in driving radiotherapy-induced anti-tumor immunity. Our study provides deeper insights into the dynamic behavior of CD8^+^ T-cells and their critical involvement in tumor control following radiotherapy.

## INTRODUCTION

Tumors must evade immune control, specifically T-cells, to progress ^1^. For instance, the density of CD8^+^ T-cells in tumors correlates with better survival in most tumor types ^2^, and mobilizing CD8^+^ T-cell-mediated immunity by immune checkpoint inhibitors has revolutionized the treatment of cancer ^3^.

Radiotherapy is a frequently used treatment for cancer; approximately half of all cancer patients will receive radiotherapy at some point ^4^. In addition to its genotoxic effect on cancer cells, radiotherapy promotes a local inflammatory response that supports tumor-specific immunity. For example, irradiated tumors have a higher density of tumor-associated effector CD8^+^ T-cells than non-irradiated tumors, and in fact, the efficacy of radiotherapy in pre-clinical models depends on CD8^+^ T-cells ^5,6^. Several studies have addressed how radiotherapy supports tumor-specific immunity, and various factors were suggested, including type I interferon (IFN) ^7–9^, increased expression of major histocompatibility complex (MHC) class I molecules and tumor-associated antigens ^10,11^, and maturation of tumor-associated dendritic cells (DCs) ^5–7^. Despite these observations, how radiotherapy influences the dynamics of different CD8^+^ T-cell subsets in the tumor remains unclear.

Tumor-infiltrating CD8^+^ T-cells display considerable inter- and intratumor heterogeneity in clinical and preclinical samples ^12,13^. Besides effector and memory cells, tumors contain CD8^+^ T-cells at different stages of differentiation including exhausted and dysfunctional exhausted subsets ^13^. Chronic exposure to antigens triggers an epigenetic differentiation program in T-cells that severely impairs their effector functions, metabolic fitness, and induces the expression of inhibitory receptors like PD-1 ^14,15^. This exhausted state is generally considered terminal. However, the proliferation of intratumoral CD8^+^ T-cells after PD-1 blockade ^16^ suggests that not all exhausted cells are terminal. In fact, a subset of virus-specific CD8^+^ T-cells co-expressing the inhibitory receptor PD-1 and the transcription factor T-cell factor 1 (TCF-1, encoded by *Tcf7*) has been shown to maintain immune responses during chronic infections ^17–19^. These so-called stem-like CD8^+^ T-cells (TCF-1^+^ PD-1^+^ TIM-3^-^, also known as progenitor exhausted or TPEX) are capable of asymmetric cell division, which allows them to both self-renew and generate effector CD8⁺ T cells (TCF-1^-^ PD-1^+^ TIM-3^+^) at the same time ^20,21^. The frequency of intratumoral stem-like CD8^+^ T-cells correlates with clinical response to PD-1 blockade ^22–24^, which has stimulated wide interest in this subset.

Additionally, tumor-draining lymph nodes have been identified as reservoirs of stem-like CD8^+^ T-cells that migrate into the tumor and play a crucial role in sustaining the anti-tumor immune response ^25,26^. However, it remains unclear how radiotherapy influences these intratumoral stem-like CD8⁺ T cells and whether they play a role in the therapeutic efficacy of radiotherapy.

In this study, we explore how intratumoral CD8^+^ T-cells respond to radiotherapy. We propose that radiotherapy drives the differentiation and proliferation of intratumoral stem-like CD8^+^ T-cells, thereby playing an important role in tumor control.

## RESULTS

### Radiotherapy leads to activation and differentiation of intratumoral CD8^+^ T-cells

To investigate how radiotherapy affects the dynamics of CD8^+^ T-cells within the tumor microenvironment, we irradiated C57BL/6 mice bearing a subcutaneous MC38 tumor with a single dose of 20 Gy (**Figure S1A**). Radiotherapy resulted in significant tumor control, both in size (**Figure S1B**) and weight (Figure S1C). Nine days after radiotherapy, we analyzed intratumoral CD8^+^ T-cells by multi-parameter spectral flow cytometry to assess radiotherapy-induced phenotypic and functional changes (**Figure S1D**). Consistent with previous findings ^5,27,28^, the number of intratumoral CD8^+^ T-cells was significantly higher after radiotherapy (**Figure S1E**). Furthermore, after gating on CD8^+^ T-cells, we observed a decreased frequency of CD62L^+^, an increased frequency of PD-1^+^ and of TIM-3^+^ cells (**Figure S1F**), and a higher frequency of granzyme B-positive (GZMB^+^) cells (**Figure S1G**). We observed an increased frequency of Ki-67^+^ intratumoral CD8^+^ T-cells (**Figure S1H**). Uniform Manifold Approximation and Projection (UMAP) visualization, combined with Rphenograph clustering, showed a distinct clustering of intratumoral CD8^+^ T-cells after radiotherapy (**Figure S1I**). The accompanying heatmap illustrates the relative expression of key markers across different clusters, enabling the simultaneous analysis of multiple markers and providing a comprehensive view of T-cell states (**Figure S1J**). In summary, these results suggest that radiotherapy induces the differentiation and proliferation of intratumoral CD8^+^ T-cells (**Figure S1K**).

### Pre-existing intratumoral CD8^+^ T-cells are sufficient for the therapeutic efficacy of radiotherapy

To study tumor-specific CD8^+^ T-cells, we used ovalbumin-expressing MC38 cells (MC38-OVA) as tumor model (**Figure S2A**). We demonstrated that the therapeutic efficacy of radiotherapy depends on CD8^+^ T-cells (**Figures S2A-D**), which is in agreement with previously published work ^5,29,30^. While the involvement of CD8^+^ T-cells in radiotherapy-dependent tumor control has been established by multiple studies, it is less clear whether *de novo* recruited or pre-existing intratumoral CD8^+^ T-cells are most important. To address this question, we injected mice with FTY720 (fingolimod) immediately before radiotherapy. FTY720 prevents the egress of lymphocytes from secondary lymphoid organs into the blood ^31^, thereby blocking *de novo* infiltration into the tumor tissue (**Figures 1A and S3A**). We will use the term “pre-existing intratumoral” for CD8^+^ T-cells that were present in the tumor before blocking new infiltration by FTY720, and the term “intratumoral” for all CD8^+^ T-cells in the tumor including pre-existing and newly infiltrated cells ^32^. We found that FTY720 did not influence the efficacy of radiotherapy (**Figures 1B and 1C**) or its effect on total and tumor-specific intratumoral CD8^+^ T-cells (**Figures 1D and 1G**). Despite FTY720 preventing *de novo* infiltration, the absolute number of intratumoral CD8^+^ T-cells was higher in irradiated tumors, suggesting that radiotherapy induces differentiation and proliferation of pre-existing intratumoral CD8^+^ T-cells. The UMAP visualization of clusters showed radiotherapy-induced differences in CD8^+^ T-cell clustering, which were not affected by FTY720 treatment. (**Figure S3B**). The heatmap illustrates the relative expression levels of markers across different clusters (**Figure S3C**). These clusters confirm that radiotherapy promotes differentiation and increases the frequency of SIINFEKL-specific CD8^+^ T-cells (**Figure S3C**), and these changes were unaffected by FTY720 (**Figure S3D**). The observation that the frequency of cluster 11 (Ki67^high^), significantly decreased after radiotherapy, regardless of FTY720 (**Figure S3D**), suggests that the initial radiotherapy-induced proliferative response diminishes nine days after radiotherapy.

**Figure 1.**
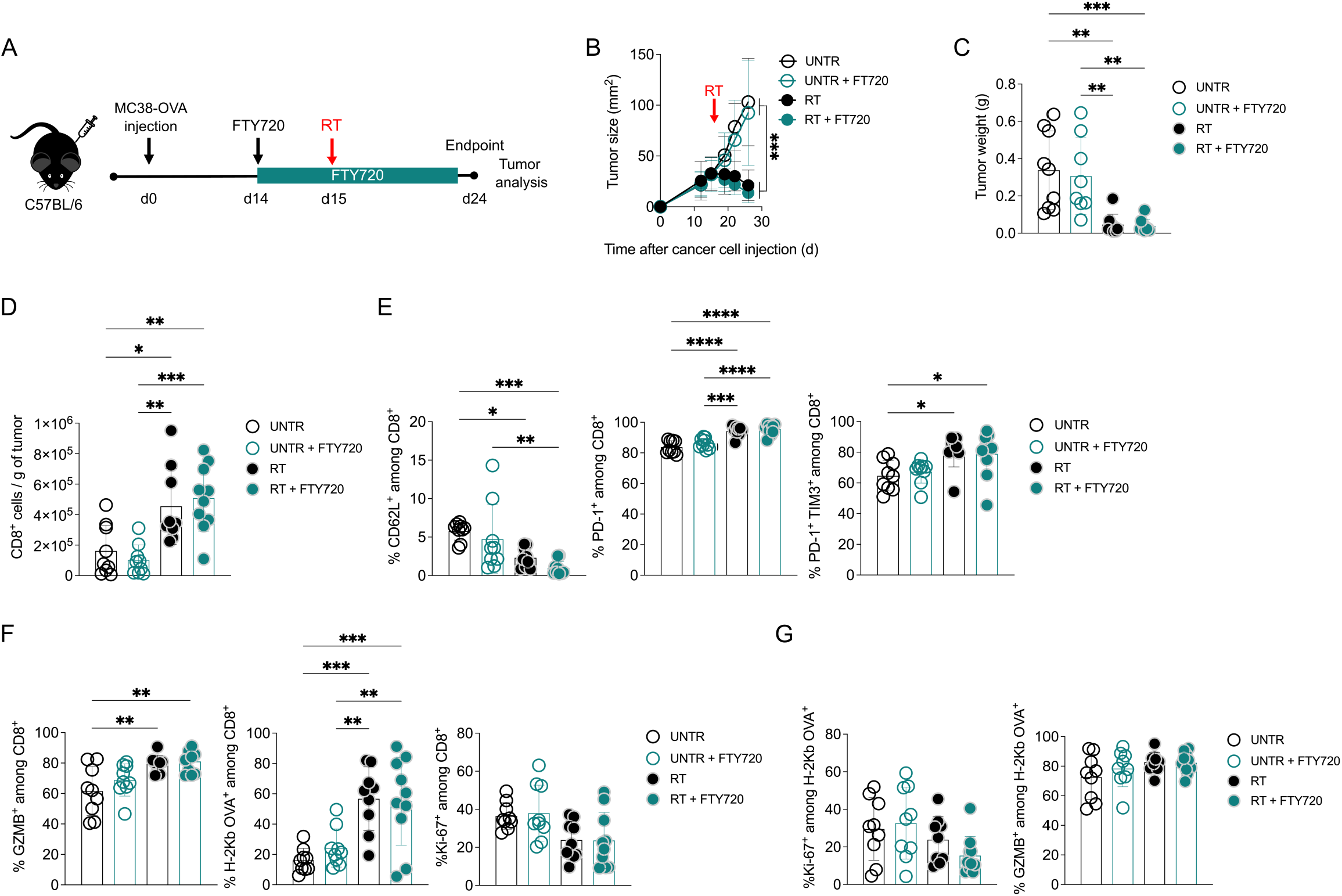
Pre-existing intratumoral CD8+ T-cells are sufficient for the therapeutic efficacy of radiotherapy. (A) Experimental setup: C57BL/6 mice (RT+FTY720 n=10, the rest n=9 per condition) were injected subcutaneously with 8x105 MC38-OVA cells in matrigel:PBS (1:2). Fifteen days later, some groups received radiotherapy (RT; 20 Gy) or not (UNTR). Starting 1 day before radiotherapy, some groups received 20 µg FTY720 per os every second day until the endpoint. Tumors were analyzed nine days after radiotherapy. (B) Tumor size (length x breadth) was measured by caliper. (C) Tumor weight (D) Number of CD8+ T-cells per gram of tumor. (E) Percentage of CD62L, PD-1, PD-1 and TIM-3 expressing CD8+ T-cells. (F) Percentage of GZMB and Ki-67 expressing CD8+ T-cells and SIINFEKL-specific CD8+ T-cells, determined by H-2Kb OVA tetramer. (G) Percentage of GZMB and Ki-67 expressing SIINFEKL-specific CD8+ T-cells. The bar indicates the mean ±SD. Groups were statistically compared using one- way ANOVA with Tukey’s multiple comparison correction. *p<0.05, **p<0.01, ***p<0.001, ****p<0.0001.

We repeated the experiment using the B16-OVA melanoma cell line (**Figure S4A**) and obtained comparable results (**Figures S4B-F**), indicating that our findings are not restricted to a single experimental model.

### Radiotherapy induces proliferation and differentiation of pre-existing intratumoral CD8^+^ T-cells

To understand the dynamics of pre-existing intratumoral CD8^+^ T-cells, we performed multiparameter single-cell mapping at different time points following radiotherapy. One (d1) or three days (d3) after radiotherapy (**Figure 2A**), no therapeutic effect was yet detectable (**Figure 2B**). Compared to untreated tumors, the number of pre-existing intratumoral CD8^+^ T-cells was significantly lower on day 1 but increased by day 3 (**Figure 2C**). Phenotypic and functional analysis revealed minimal changes, except for a significant decrease in Ki-67^+^ CD8^+^ T-cells on day 1, suggesting an increased sensitivity of proliferating CD8^+^ T-cells to radiotherapy-induced cell death (**Figures S5A and S5B**). UMAP and clustering analysis supported these findings, showing no significant clustering differences apart from a reduced frequency of the Ki-67^high^ cluster (**Figures S5C-E**).

**Figure 2.**
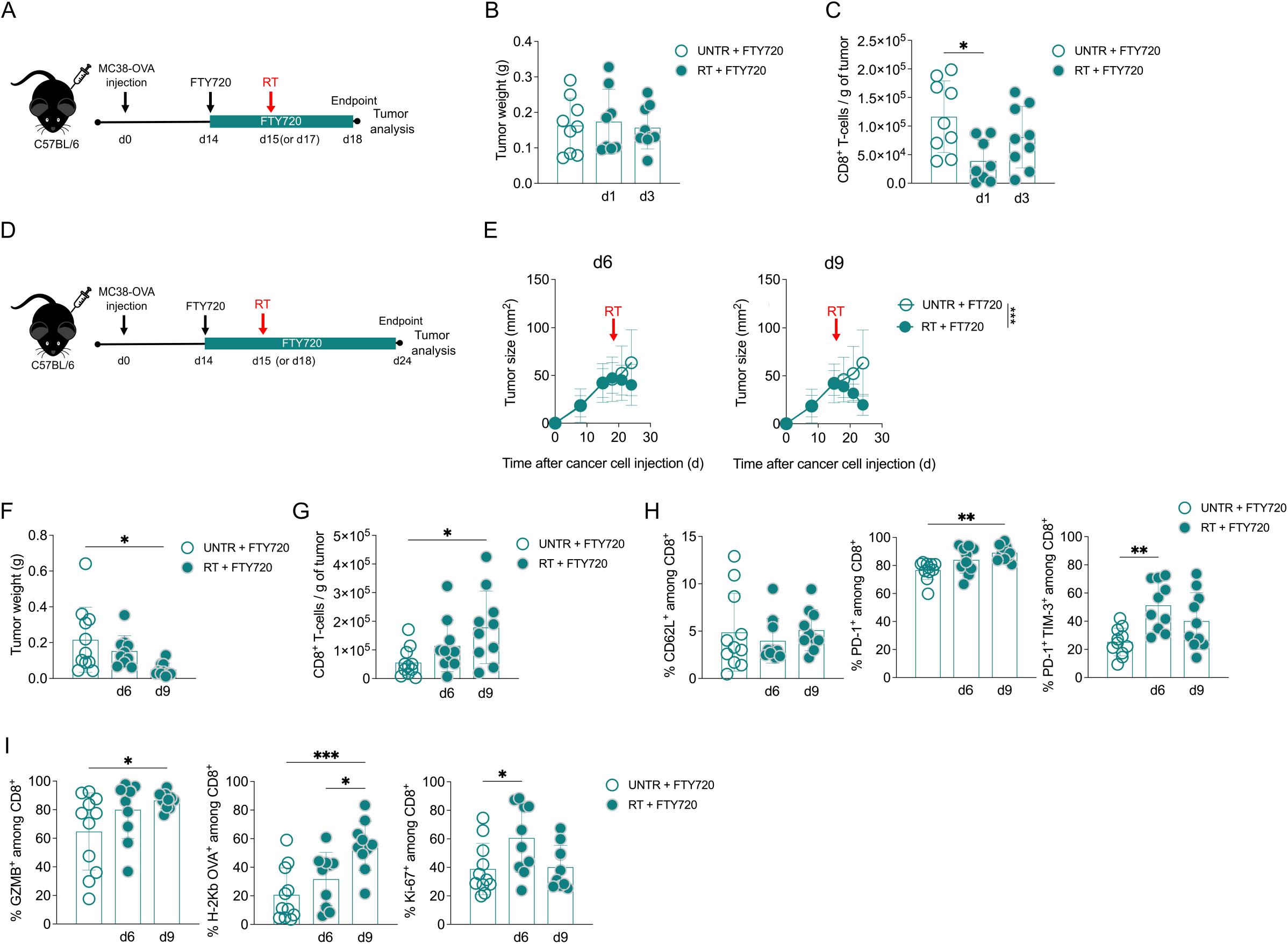
The initial loss of pre-existing intratumoral CD8+ T-cells is followed by proliferation after radiotherapy. (A) Experimental setup: C57BL/6 mice (n=9 per condition) were injected subcutaneously with 8x105 MC38-OVA cells in matrigel:PBS (1:2). Fifteen (d3) or seventeen (d1) days later, some groups received radiotherapy (RT; 20 Gy) or not (UNTR). Starting 1 day before radiotherapy, all groups received 20 µg FTY720 per os every second day until the endpoint. Tumors were analyzed one or three days after radiotherapy. (B) Tumor weight. (C) Number of CD8+ T-cells per gram of tumor. (D) Experimental setup: C57BL/6 mice (UNTR n=11, the rest n=10 per condition) were injected subcutaneously with 8x105 MC38-OVA cells in matrigel:PBS (1:2). Fifteen (d9) or eighteen (d6) days later, some groups received radiotherapy (RT; 20 Gy) or not (UNTR). Starting 1 day before radiotherapy, all groups received 20 µg FTY720 per os every second day until the endpoint. Tumors were analyzed six or nine days after radiotherapy. (E) Tumor size (length x breadth) was measured by caliper. (F) Tumor weight (G) Number of CD8+ T-cells per gram of tumor. (H) Percentage of CD62L, PD-1, PD-1 and TIM-3 expressing CD8+ T-cells. (I) Percentage of GZMB and Ki-67 expressing CD8+ T-cells and SIINFEKL-specific CD8+ T-cells, determined by H-2Kb OVA tetramer. The bar indicates the mean ±SD. Groups were statistically compared using one-way ANOVA with Tukey’s multiple comparison correction. *p<0.05, **p<0.01, ***p<0.001.

On days six (d6) and nine (d9) after radiotherapy, we observed progressive tumor reduction in both size and weight, indicating ongoing tumor control (**Figures 2D-F**). Correspondingly, the number of pre-existing intratumoral CD8^+^ T-cells continued to increase, suggesting that CD8^+^ T-cell proliferation remains active during this period (**Figure 2G**). Although no significant changes were observed in the frequency of early differentiated CD62L^+^ CD8^+^ T-cells, we observed an increase in differentiated CD8^+^ T-cells expressing PD-1 and TIM-3 at both time points (**Figure 2H**). Additionally, the frequency of GZMB^+^ and SIINFEKL-specific CD8^+^ T-cells continued to increase between day 6 and 9 (**Figure 2I**). Consistent with earlier observations (**Figure 1F**), the frequency of Ki-67^+^ CD8^+^ T-cells significantly increased on day 6, followed by a decrease on day 9, indicating that radiotherapy induces a transient proliferative response in pre-existing intratumoral CD8^+^ T-cells. UMAP analysis further reflected these dynamic changes (**Figures S6A-C**). The frequency of clusters with high Ki-67 expression, such as cluster 4, peaked on day 6 and declined by day 9, corroborating our findings (**Figure S6C**).

### Radiotherapy induces proliferation and differentiation of pre-existing intratumoral stem-like CD8^+^ T-cells

To gain a deeper understanding of the radiotherapy-induced changes in intratumoral CD8^+^ T-cells, we performed single-cell RNA sequencing on sorted pre-existing and intratumoral CD8^+^ T-cells six days after radiotherapy (**Figure 3A**). We used the ProjecTILs package ^33^ to annotate the intratumoral CD8^+^ T-cells and identified five distinct CD8^+^ T-cell phenotypes: naïve-like, early active, stem-like, effector/memory, and exhausted (**Figure 3B**). These differentiation states were correctly annotated, as confirmed by radar plots (**Figure S7A**) and violin plots (**Figure S7B**). Radiotherapy reduced the frequency of naïve-like intratumoral CD8^+^ T-cells, regardless of FTY720 treatment (**Figures 3C and S8**). Without FTY720, there was an increase in early activated and effector/memory CD8^+^ T-cell states, while blocking new infiltration led to more exhausted CD8^+^ T-cells. This suggests that pre-existing CD8^+^ T-cells are more differentiated and exhausted compared to intratumoral CD8^+^ T-cells, which likely include newly infiltrated cells in earlier differentiation states (**Figures 3C and S8**).

**Figure 3.**
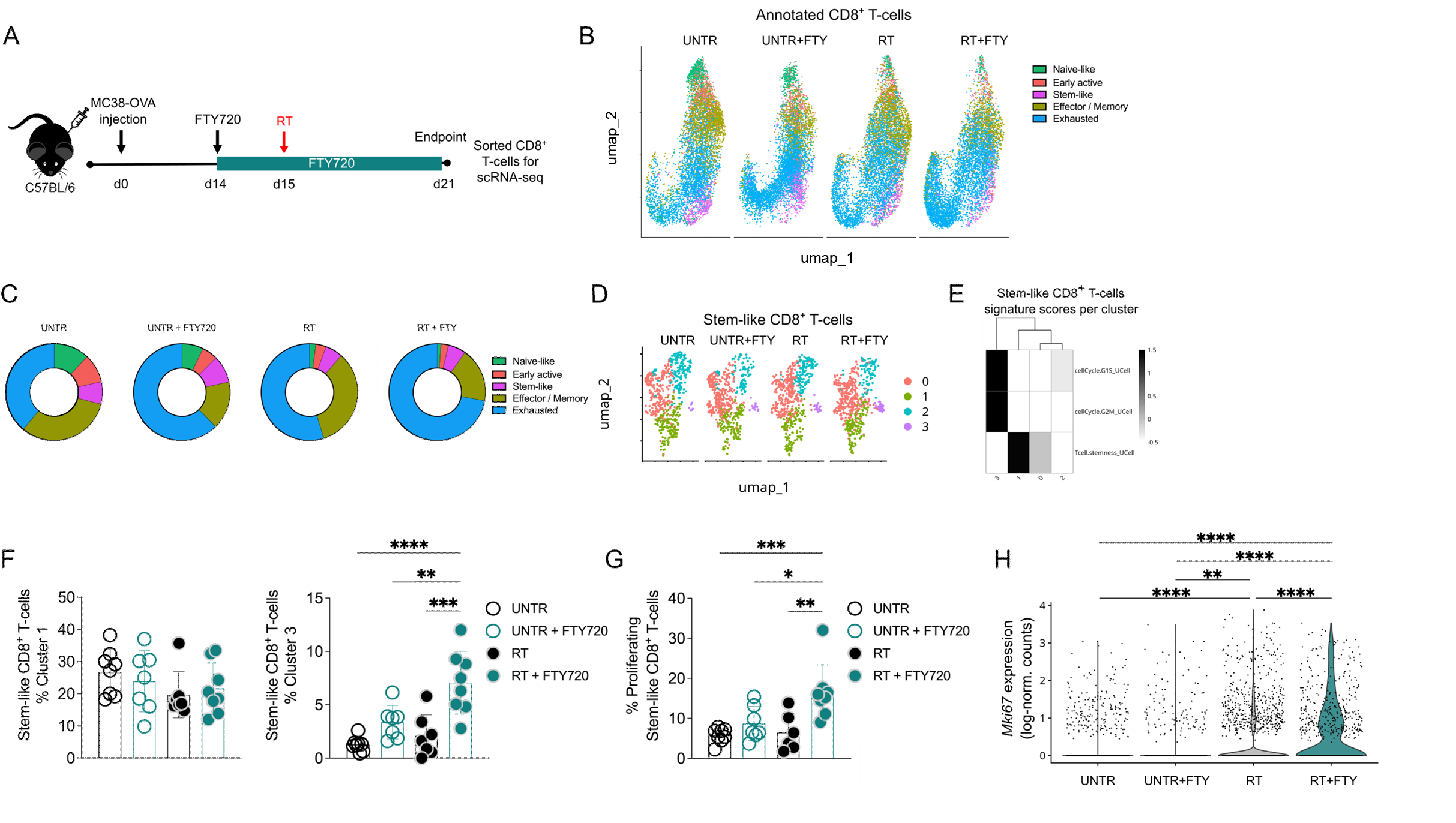
Radiotherapy promotes the proliferation of pre-existing intratumoral stem-like CD8+ T-cells. (A) Experimental setup: C57BL/6 mice (UNTR+FTY720 n=7, the rest n=8 per condition) were injected subcutaneously with 8x105 MC38-OVA cells in matrigel:PBS (1:2). Fifteen days later, some groups received radiotherapy (RT; 20 Gy) or not (UNTR). Starting 1 day before radiotherapy, all groups received 20 µg FTY720 per os every second day until the endpoint. Six days after radiotherapy, tumor samples were hashed and CD8+ T-cells were sorted using flow cytometry. Samples were prepared using the BD Rhapsody platform and sequenced on the Illumina NovaSeq X. (B) UMAP representation of annotated CD8+ T-cell subsets, determined by ProjectTILs package. (C) Pie chart showing frequencies of CD8+ T-cell subsets per condition. (D) UMAP representation of stem-like CD8+ T-cell clusters. (E) Heatmap illustrates signature scores (determined by UCell package) across different stem-like CD8+ T-cell clusters. (F) Percentage of UMAP clusters 1 and 3. (G) Percentage of proliferating stem-like CD8+ T-cells (defined by a G1S or G2M score >0.1), determined by UCell package. (H) Mki67 expression shown as log-normalized counts. The bar indicates the mean ±SD. Groups were statistically compared using one-way ANOVA with Tukey’s multiple comparison correction. *p<0.05, **p<0.01, ***p<0.001, ****p<0.0001.

Further analysis of exhausted CD8^+^ T-cells revealed five distinct clusters (**Figure S9A**). We used single-cell gene signature scoring using the UCell package to characterize these clusters ^34^. Among the two most abundant clusters, cluster 0 exhibited pronounced signatures of exhaustion and interferon response, while cluster 1 showed a strong proliferation signature (**Figures S9B and S9C**). After radiotherapy, cluster 0 was significantly enriched in total intratumoral CD8^+^ T-cells, supporting our observation that radiotherapy promotes differentiation of pre-existing intratumoral CD8^+^ T-cells. Further, pre-existing intratumoral CD8^+^ T-cells were significantly enriched in cluster 1, indicating a robust radiotherapy-induced proliferative response (**Figure S9D**). To further validate these findings, we applied a threshold to the G1S and G2M UCell scores for all exhausted CD8^+^ T-cells, classifying cells with scores above this threshold as proliferating. We found that the frequency of proliferating exhausted CD8^+^ T-cells was indeed higher in pre-existing intratumoral CD8^+^ T-cells after radiotherapy (**Figure S9E**). Additionally, there was a significant increase in *Mki67* expression after radiotherapy, especially in pre-existing CD8^+^ T-cells (**Figure S9F**).

ProjecTILs package effectively identified stem-like CD8^+^ T-cells, a subset that has been challenging to detect using flow cytometry in our tumor models (**Figures 3B and 3C**).

Stem-like CD8^+^ T-cells (defined as TCF1^+^PD-1^+^) are essential for the response to anti-PD-1 therapy ^20–24^ and are thought to be self-renewing precursors of effector T-cells. Further analysis revealed four distinct clusters of intratumoral stem-like CD8^+^ T-cells (**Figure 3D**). Using UCell analysis, we found that cluster 1 had the highest stemness score, closely resembling the previously characterized CD62L^+^ stem-like population ^35^. Cluster 3 showed a strong proliferation signature, displaying high expression of genes associated with the G1S and G2M phases (**Figure 3E**). While the frequency of stem-like CD8^+^ T-cells in cluster 1 remained unchanged, the number of cells in cluster 3 significantly increased among pre-existing intratumoral stem-like CD8^+^ T-cells after radiotherapy (**Figure 3F**). To validate these findings, we applied the same threshold to the G1S and G2M UCell scores for all stem-like CD8^+^ T-cells. This analysis confirmed that the frequency of proliferating stem-like CD8^+^ T-cells was indeed higher in pre-existing intratumoral CD8^+^ T-cells after radiotherapy (**Figure 3G**). Moreover, *Mki67* expression was significantly higher after radiotherapy, particularly in pre-existing CD8^+^ T-cells (**Figure 3H**). Together, these data suggest that pre-existing intratumoral stem-like CD8^+^ T-cells proliferate in response to radiotherapy.

After observing dynamic changes in subsets of intratumoral CD8^+^ T-cells following radiotherapy, we aimed to identify the differentiation trajectory of these cells using the Monocle 3 package ^36,37^. We selected a root node from the TCF-1^high^, CD62L^high^, and PD-1^low^ region (**Figure S10A**). The inferred trajectory was plotted on a UMAP, both for annotated clusters (left) and pseudotime (right), further confirming that exhausted CD8^+^ T-cells represent the most differentiated cell state compared to the naïve-like state (**Figure S10B**). The expression of key differentiation and activation markers along the pseudotime trajectory confirmed that pseudotime accurately reflects the biological differentiation states of CD8^+^ T-cells. For example, CD62L and TCF-1 expression progressively decrease over pseudotime, while PD-1 and TIM-3 increase in expression (**Figure S10C**). Trajectory inference also identified three gray nodes as different outcomes of the trajectory within the stem-like CD8^+^ T-cell population, which are directly linked to the exhausted and effector/memory states. This suggests that intratumoral stem-like CD8^+^ T-cells may indeed be precursors of differentiated CD8^+^ T-cells in response to radiotherapy.

### TCF-1^+^ cells are essential for the efficacy of radiotherapy

To evaluate the importance of TCF-1^+^PD-1^+^ stem-like CD8^+^ T-cells, we used the *Tcf7*^DTR-GFP^ mouse model to selectively deplete TCF-1^+^ cells using diphtheria toxin (DTX) ^20^. To avoid systemic depletion of TCF-1^+^ cells, Thy1.1^+^ *Tcf7*^DTR-GFP^ donor bone marrow was transplanted into lethally irradiated Thy1.2^+^ C57BL/6 recipient mice. The remaining host T-cells ^38,39^ (**Figure S11A**) were depleted by anti-Thy1.2 antibodies (**Figure S11B**). After reconstitution, mice received radiotherapy (or not) and DTX (or not), and all groups received FTY720 (**Figures 4A and S11C**). At the endpoint, we confirmed TCF-1^+^ cell depletion by assessing GFP^+^ expression in CD8^+^ T-cells from the spleen (**Figure S11D**) and the tumor (**Figure S11E**).

**Figure 4.**
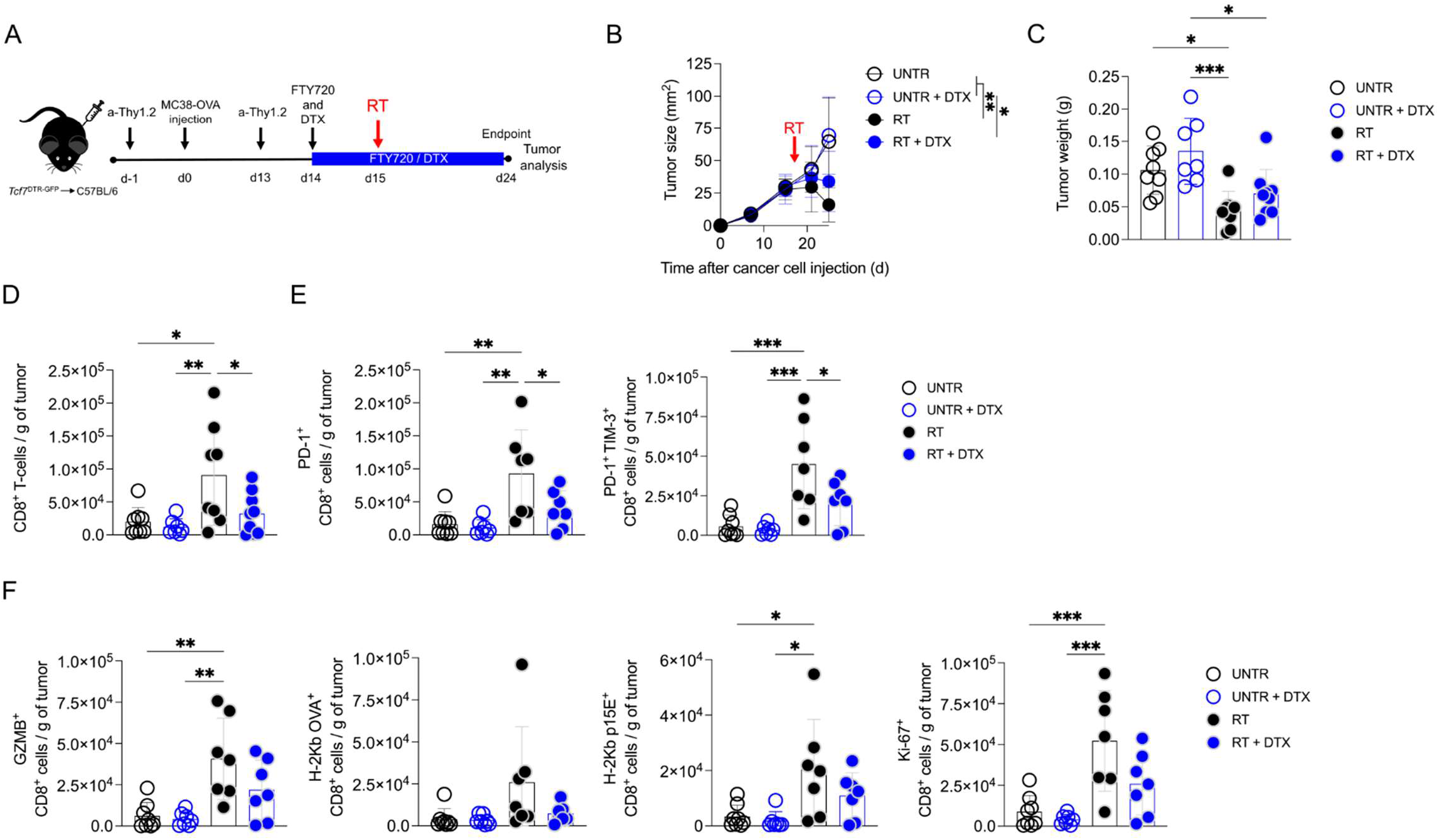
Depletion of TCF-1+ cells impairs the efficacy of radiotherapy. (A) Experimental setup: Tcf7DTR-GFP→C57BL/6 mice (UNTR, RT n=8, UNTR+DTX n=7, RT+DTX n=9) bone marrow transplant was performed as described (22685313). After 8 weeks, the mice were injected subcutaneously with 8x105 MC38-OVA-GFP cells in matrigel:PBS (1:2). One day before tumor injection and one day before radiotherapy, the remaining host T-cells were depleted by a double intraperitoneal injection of 500 µg of anti-Thy1.2 (30H12). Fifteen days after tumor injection, some groups received radiotherapy (RT; 20 Gy) or not (UNTR). Starting 1 day before radiotherapy, all groups received 20 µg FTY720 per os every second day, some groups received 250 ng diphtheria toxin (DTX) intraperitoneal injection twice per week, whereas the other groups received PBS until the endpoint. Tumors were analyzed nine days after radiotherapy. (B) Tumor size (length x breadth) was measured by caliper. (C) Tumor weight (D) Number of CD8+ T-cells per gram of tumor. (E) Number of PD-1 and PD-1 and TIM-3 expressing CD8+ T-cells per gram of tumor. (F) Number of GZMB and Ki-67 expressing CD8+ T-cells, SIINFEKL-specific (determined by H-2Kb OVA tetramer) and p15-specific (determined by H-2Kb p15E tetramer) CD8+ T-cells per gram of tumor. Each symbol represents one mouse. The bar indicates the mean ±SD. Groups were statistically compared using one-way ANOVA with Tukey’s multiple comparison correction. *p<0.05, **p<0.01, ***p<0.001.

Radiotherapy decreased tumor progression and weight, and this effect was slightly diminished when TCF-1^+^ cells were depleted (**Figures 4B and 4C**). We also observed the expected increase in CD8^+^ T-cell counts after radiotherapy, which was significantly reduced after TCF-1^+^ cell depletion (**Figure 4D**). In line with our previous observations, the number of differentiated (**Figure 4E**), cytotoxic, SIINFEKL-specific, p15E-specific ^40^, and proliferating CD8^+^ T-cells was significantly higher after radiotherapy, but this increase was limited after TCF-1^+^ cell depletion (**Figure 4F**). These findings suggest that TCF-1^+^ cells are precursors of proliferating and differentiating T-cells in the tumor. UMAP analysis confirmed that the radiotherapy-induced changes in pre-existing intratumoral CD8^+^ T-cells depend on TCF-1^+^ cells (**Figure S12**). Furthermore, the increase in the number of SIINFEKL-specific (cluster 4) and Ki-67^high^ (cluster 6) CD8^+^ T-cells depends on TCF-1^+^ cells (**Figures S12B and S12C**).

To corroborate these findings and examine the role of stem-like CD8^+^ T-cells in long-term radiotherapy-induced tumor control, we conducted the TCF-1^+^ depletion experiment with a focus on survival (**Figure S13A**). We generated Thy1.1^+^ *Tcf7*^DTR-GFP^ → Thy1.2^+^ *Tcrb*^-/-^ ^41^ bone marrow chimeras, which circumvented the need to deplete host T-cells (**Figures S13B**). We treated all groups with FTY720 to block new T-cell infiltration (**Figure S13C**) and validated TCF-1^+^ cell depletion by analyzing GFP^+^ expression in CD8^+^ T-cells from the spleen at the time of sacrifice (**Figure S13D**). We monitored the tumors for 50 days and observed a significant tumor control following radiotherapy, but this effect was reduced after TCF-1^+^ cell depletion (**Figure S13E**).

Although no significant change in survival was observed due to biological variation (**Figure S13F**), we found a decreased proportion of mice responding to radiotherapy after TCF-1^+^ depletion (**Figure S13G**). These observations suggest that TCF-1^+^ cells play a crucial role in radiotherapy-induced immunological and clinical response.

### Stem-like CD8^+^ T-cells differentiate and proliferate after radiotherapy

To investigate whether stem-like CD8^+^ T-cells are the precursors of radiotherapy-induced effector CD8^+^ T-cells, we sorted intratumoral TCF-1^-^PD-1^+^ (effector) and TCF- 1^+^PD-1^+^ (stem-like) CD8^+^ T-cells (**Figure S14**) and adoptively transferred 3,000 sorted CD8^+^ T-cells (CD45.1) into tumor-bearing *Tcrb*^-/-^ mice. Mice received radiotherapy one day later, and CD8^+^ T-cells from the tumor and tumor-draining lymph nodes were analyzed nine days after radiotherapy (**Figure 5A**). Tumor size and weight did not significantly differ between mice injected with stem-like or effector CD8^+^ T-cells (**Figures 5B and 5C**). However, six days after radiotherapy we found a significantly higher frequency of CD45.1^+^ CD8^+^ T-cells in the blood of mice injected with stem-like CD8^+^ T-cells (**Figure 5D**). Similarly, mice injected with stem-like CD8^+^ T-cells showed significantly higher counts of CD8^+^ T-cells in both the tumor (**Figure 5E**) and the tumor-draining lymph nodes (**Figure 5F**). UMAP analysis of stem-like CD8^+^ T-cell recipient mice revealed distinct clustering and phenotypic differences in CD8^+^ T-cells collected from tumors and lymph nodes (**Figure S15A and S15B).** In the lymph nodes, most CD8^+^ T-cells maintained a less differentiated phenotype (TCF-1^high^ PD-1^low^), whereas intratumoral CD8^+^ T-cells showed a more differentiated phenotype (TCF-1^low^ PD-1^high^) (**Figures S15A and S15B**). Early differentiated clusters, such as cluster 1 (PD-1^low^TIM- 3^low^), were significantly more frequent in the lymph nodes, while more differentiated clusters, like cluster 4 (PD-1^high^TIM-3^high^), were significantly higher in the tumor (**Figures S15C and S15D**). These results suggest that stem-like CD8^+^ T-cells preserve their stemness in the lymph nodes but differentiate and proliferate upon reaching the tumor microenvironment.

**Figure 5.**
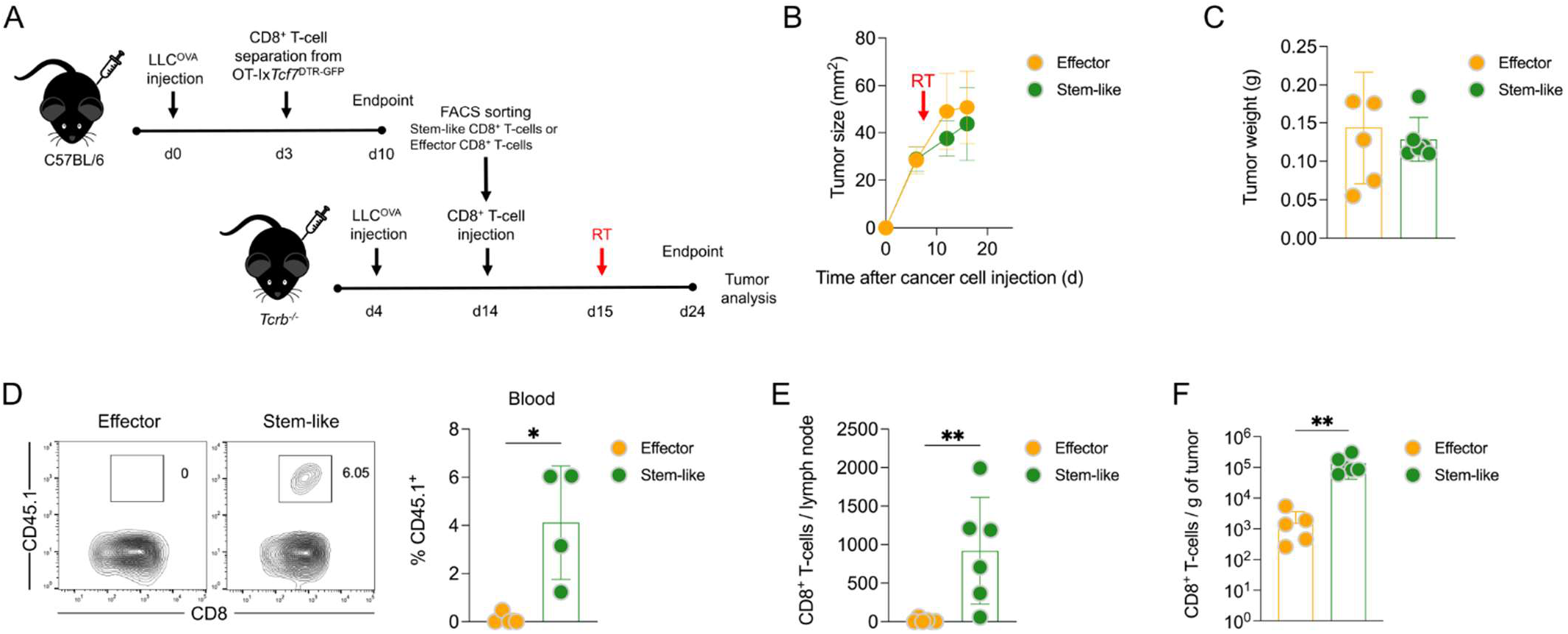
Stem-like CD8+ T-cells differentiate and proliferate after radiotherapy. (A) Experimental setup: C57BL/6 mice (n=15) were injected subcutaneously with 3x105 LLC-OVA cells in matrigel:PBS (1:2). Three days later, all mice received an intravenous injection of magnetically enriched OT-I x Tcf7DTR-GFP (CD45.1) CD8+ T-cells (105 cells per mouse). Seven days later, tumors were processed, and stem-like (TCF-1+PD-1+) and effector (TCF-1−PD-1+) CD8+ T-cells were sorted by flow cytometry. These cells were then intravenously injected into LLC-OVA-bearing Tcrb-/- mice (3000 cells per mouse). One day later, all mice (n=6 per condition) received radiotherapy (RT; 20 Gy). Tumors and tumor-draining lymph nodes were analyzed nine days after radiotherapy. (B) Tumor size (length x breadth) was measured by caliper. (C) Tumor weight. (D) Presence of adoptively transferred cells was evaluated by measuring the percentage of CD45.1+ cells among CD45+ cells in the blood six days after radiotherapy. (E-F) Number of CD8+ T-cells per lymph node (E) and per gram of tumor (F). Each symbol represents one mouse. The bar indicates the mean ±SD. Groups were statistically compared using the Mann-Whitney U test. *p<0.05, **p<0.01.

To further validate these findings and confirm that radiotherapy is essential for the stem-like CD8^+^ T-cell response, we conducted a similar experiment including untreated controls. This time, *Tcrb*^-/-^ mice received 1,000 sorted cells (**Figure S16A**). Unlike in the previous experiment, we detected few transferred cells in the blood six days after radiotherapy, suggesting that 1,000 cells might be insufficient for detection in circulation (**Figure S16B**). We observed radiotherapy-induced tumor control independent of whether stem-like or effector CD8^+^ T-cells were transferred (**Figures S16C and S16D**). However, despite the lower cell count, radiotherapy significantly increased CD8^+^ T-cell count in the lymph nodes (**Figure S16E**) and the tumor (**Figure S16F**) in mice that received stem-like CD8^+^ T-cells. Together, these findings suggest that stem-like CD8^+^ T-cells proliferate and differentiate into effectors in response to radiotherapy, and as such play a crucial role in radiotherapy-induced tumor control by serving as precursors to effector CD8^+^ T-cells.

## DISCUSSION

Radiotherapy enhances the infiltration of CD8^+^ T-cells and other immune cells into the tumor microenvironment, promoting their differentiation and activation ^5–7,27,28,30^.

However, the temporal dynamics of these changes, particularly in pre-existing intratumoral CD8^+^ T-cells, have not been fully explored. We found that radiotherapy-induced tumor control was independent of *de novo* recruited CD8^+^ T-cells. This finding is consistent with a previous study ^32^ and implies two key points; some pre-existing intratumoral CD8^+^ T-cells are radioresistant, and radiotherapy promotes the proliferation and differentiation of these radioresistant CD8^+^ T-cells. One day after radiotherapy, we observed a decrease in the number of pre-existing intratumoral CD8^+^ T-cells, particularly of the proliferating fraction. Although intratumoral CD8^+^ T-cells are generally considered more radioresistant compared to other tissues, such as the lymphoid organs ^32^, it has been reported that proliferating CD8^+^ T-cells are more sensitive to radiotherapy-induced cell death ^42^. Supported by previous studies ^32,43^, some pre-existing intratumoral CD8^+^ T-cells survive after radiotherapy. The surviving cells demonstrated a proliferative response, peaking six days after radiotherapy. We depleted TCF-1^+^ precursors to assess their role and found that, in their absence, radiotherapy-induced tumor control and CD8^+^ T-cell expansion were significantly impaired but not completely abrogated. This suggests that TCF-1^+^ cells, including stem-like CD8^+^ T-cells, are crucial for radiotherapy response, though TCF-1^-^ CD8^+^ T-cells also contribute to tumor control. This fits with previous work showing that differentiated CD8^+^ T-cells contributed to the response to PD-1 blockade ^44–46^. Thus, while different subsets of CD8^+^ T-cells contribute to the efficacy of radiotherapy, we demonstrated that the small population of intratumoral stem-like CD8^+^ T-cells is indispensable for an effective response. The adoptive transfer of intratumoral stem-like CD8^+^ T-cells further validated their importance for radiotherapy response. These transferred stem-like CD8^+^ T-cells differentiated and proliferated in response to radiotherapy within the tumor.

Taken together, by showing that intratumoral stem-like CD8^+^ T-cells are precursors to radiotherapy-induced protective effector cells, we identified the mechanism explaining why irradiated tumors contain more effector CD8^+^ T-cells.

We showed that pre-existing intratumoral CD8^+^ T-cells can drive an effective radiotherapy response during the initial phase, resulting in the differentiation of CD8^+^ T-cells in the tumor. For example, adoptively transferred stem-like CD8^+^ T-cells preserved their stemness in the tumor-draining lymph node, whereas in response to radiotherapy they rapidly differentiated into effector cells in the tumor. This finding aligns with previous work showing that lymph nodes are reservoirs for stem-like CD8^+^ T-cells that migrate into tumors, differentiate locally, and help restrict tumor progression ^25,26,47^. Our sequencing results confirm this process, showing that intratumoral CD8^+^ T-cells acquired a more differentiated phenotype after radiotherapy. The differentiation was even more pronounced when *de novo* infiltration was blocked. This suggests that while pre-existing intratumoral CD8^+^ T-cells are sufficient for immediate responses, continued recruitment from lymph nodes may be essential for long-term anti-tumor response.

PD-1/PD-L1 blockade synergizes with radiotherapy as shown in preclinical models ^27,43^; therefore, this combination is currently explored in cancer patients ^48–52^ with promising results ^53^. Stem-like CD8^+^ T-cells (TCF1^+^ PD-1^+^) are crucial for the response to anti-PD- 1 therapy by serving as a reservoir for effector T-cells ^20–24^. Our data suggest that stem-like CD8^+^ T-cells play a similar role in the response to radiotherapy, potentially explaining the synergy seen after combining both treatments ^54^.

In conclusion, we demonstrated that radiotherapy promotes the proliferation and differentiation of pre-existing intratumoral CD8^+^ T-cells, including stem-like CD8^+^ T-cells.

This process is required and sufficient for therapeutic efficacy during the initial phase, while sustained anti-tumor response may require the recruitment of stem-like CD8^+^ T-cells from the tumor-draining lymph nodes.

## Limitation of the study

One limitation of this study is the use of a single 20 Gy radiotherapy dose. While similar transcriptional changes and CD8^+^ T-cell activation have been observed between 5 Gy, 10 Gy, and 20 Gy doses ^5,6^, the effects of varying radiation doses on our findings still need to be explored. Additionally, single high-dose radiotherapy may not fully capture the effects of fractionated radiotherapy commonly used in clinical settings. Fractionation is known to influence immune responses, including CD8^+^ T-cell dynamics, even when the biologically effective dose is equivalent ^29,30,43,55,56^. Inherently, it is also challenging to compare single-dose and fractionated radiotherapy because the timing of the treatments differs, making it difficult to align observations and assess immune responses consistently. Future studies are needed to determine how much of our findings also apply to fractionated treatments. This study exclusively uses murine tumor models, and since the tumor microenvironment and immune interactions in humans may differ, the clinical applicability of our observations requires further investigation.

Although we validated some of our observations in an orthotopic B16-OVA model, the majority of the study utilized non-orthotopic, subcutaneous models (LLC and MC38). While subcutaneous models were practical for tumor localization and measurement, they do not fully replicate the physiological conditions of orthotopic tumors ^57,58^. Further studies are needed to confirm how much of these observations extend to other orthotopic models.

## Acknowledgments

This work was supported by the University Research Priority Program “Translational Cancer Research” (University of Zurich; MvdB), the University of Zurich Forschungkredit (HK), the Hartmann-Müller Foundation Zurich (HK, MvdB), the Swiss National Science Foundation (310030_208145 MvdB), the Stiftung für Krebsbekämpfung Zurich (MvdB), the Zurich Cancer League (HK), the Stiftung für medizinische Forschung an der Medizinischen Fakultät der UZH (HK), the Monique Dornonville de la Court Foundation (MvdB). The authors thank the personnel of the Laboratory Animal Services Center (LASC, University of Zurich), particularly Jan Jerzyk, for expert animal care. We thank Tatiane Gorski (Cytometry Facility, University of Zurich) and Hubert Rehrauer (Functional Genomic Center Zurich) for help with scRNA-sequencing.

## Author contributions

HK and MvdB conceived the experiments and wrote the manuscript; HK, MH, NR and NEB performed the experiments. PP and VC performed sorting. HK, MH, MN and CB analyzed the data. MvdB and HK secured funding; All the authors reviewed the results and approved the final manuscript.

## Ethical approval statement

Mouse experiments were performed according to Swiss cantonal and federal regulations on animal protection and approved by the cantonal veterinary office of Zurich under license numbers 38/2021 (33443) and 170/2021 (34178).

## STAR METHODS

### RESOURCE AVAILABILITY

#### Lead contact

Further information and requests for resources and reagents should be directed to and will be fulfilled by the lead contact, Maries van den Broek.

#### Materials availability

The materials generated for this study can be provided upon reasonable request.

#### Data availability

All relevant data supporting the findings of this study are provided in the Methods section. Detailed information on mouse models, reagents, and software used in the research is also described in the Methods.

### EXPERIMENTAL MODEL DETAILS

#### Mice

C57BL/6NRj mice were purchased from Janvier Labs. *Tcf7*^DTR-GFP^ mice ^22^ were provided by Werner Held (University of Lausanne). IgHaThy1.1 (B6.Cg-Gpi1a Thy1a Igha) ^59–61^ mice were provided by Nicole Joller (University of Zurich). *Tcf7*^DTR-GFP^ x IgHaThy1.1 and *Tcf7*^DTR-GFP^ x OT-I ^62^ x Ly5.1 mice were bred in-house. *Tcrb*^-/-^ (B6.129P2-*Tcrb^tm1Mom^*/J) ^41^ mice were provided by Annette Oxenius (ETH Zurich, Switzerland). All strains have a C57BL/6 background. Breeding and experiments were performed under specific pathogen-free (SPF) conditions in facilities of the Laboratory Animal Services Center (LASC) at the University of Zurich. Mice had access to food and water containing 0.5 ppm ClO2 *ad libitum* and were kept in a 12-hour light/dark cycle. Experiments were performed using female mice aged 8-12 weeks unless otherwise stated. All procedures were approved by the Cantonal Veterinary Office Zurich under licenses ZH170/2021 and ZH38/2021.

#### Cell lines and cell culture

MC38 colon adenocarcinoma was obtained from Kerafast, LLC and B16F10 melanoma cells were obtained from ATCC. MC38-OVA-GFP^dim^ cells were provided by Mark J. Smyth ^63^,QIMR Berghofer Medical Research Institute, Brisbane, Australia). B16F10 and LLC cells were lentivirally transduced to express OVA using the pHR OVA/p2a/mCherry-CaaX plasmid (Addgene #113030). Six days after transduction, cells were sorted for live (Zombie NIR^−^) mCherry^+^ cells. Cancer cells were cultured in DMEM (Gibco) supplemented with 10% fetal bovine serum (Gibco), 100 U/mL penicillin, 100 μg/mL streptomycin (Sigma), and 2 mM L-glutamine (Gibco) at 37°C in a humidified atmosphere with 5% CO2. Cell lines were tested negative for *Mycoplasma* ssp. by PCR. Cells were also tested negative for 18 additional mouse pathogens using the IMPACT II Test (IDEXX Bioanalytics).

### METHOD DETAILS

#### *In vivo* experiments

*Cancer cell injection*. Cultured cancer cells were harvested and suspended in a 2:1 mixture of PBS and Matrigel Basement Membrane Matrix (Corning). The number of injected cells is specified in the figure legends. Hundred µl of the cell suspension was injected subcutaneously (s.c.) into the shaved right flank. For survival studies, mice were euthanized at the humane endpoint, i.e., when tumor size reached 150 mm² or the tumor developed an invagination > 30% of its surface.

*Depletion of cells*. To deplete T-cells, mice received two injections of 500 µg of CD4- depleting antibody (clone GK1.5; Rat IgG2a) or CD8-depleting antibody (clone YTS169.4; Rat IgG2a) intraperitoneally (i.p.) at days −1 and +12 relative to radiotherapy. T-cell depletion was confirmed by flow cytometric analysis of blood taken two days after the first injection.

To deplete the remaining host T-cells from Thy1.1^+^ *Tcf7*^DTR-GFP^ → Thy1.2^+^ C57BL/6 bone marrow chimeras, 500 µg of Thy1.2-depleting antibody (clone 30H12; Rat IgG2b) was injected i.p. on day −1 before tumor cell injection and on day −1 relative to radiotherapy. Depletion was verified by flow cytometry analysis of blood samples.

To deplete TCF-1 expressing cells in *Tcf7*^DTR-GFP^ mice, 250 ng of diphtheria toxin (DTX) was injected i.p. twice a week, starting one day before radiotherapy. Depletion was monitored by analyzing blood samples during the experiment and confirmed at the endpoint in the spleen and tumor tissues using flow cytometry.

*FTY720 administration*. FTY720 (Sigma-Aldrich) was administered *per os* by micropipette-guided drug administration (MDA), as previously described ^64^. Mice received 20 µg FTY720 diluted in 80 µL condensed milk (diluted 1:2 with sterile water). The reduction of circulating lymphocytes was confirmed by analyzing blood samples using flow cytometry.

*Adoptive transfer of T-cells*. For adoptive cell transfer, CD8^+^ T-cells were isolated from the spleens of age-matched OT-I mice using the EasySep Mouse CD8^+^ T-Cell Isolation Kit (STEMCELL Technologies) according to the manufacturer’s instructions. The purity of the enriched CD8^+^ T-cells (> 95%) was verified by flow cytometry. Purified CD8^+^ T-cells were injected into recipient mice via the lateral tail vein in 200 μL of sterile PBS. The number of transferred cells is specified in the figure legends.

*Radiotherapy*: Tumor-bearing mice were randomized based on tumor size immediately before radiotherapy. Radiotherapy was performed as described ^6^. Briefly, mice were anesthetized by i.p. injection of 50 mg/kg ketamine and 10 mg/kg xylazine. Vitamin A eye ointment was applied to the eyes to prevent dryness during the procedure. Mice were secured in a lead cage to ensure localized irradiation of the tumor with a single dose of 20 Gy using an RS-2000 irradiation unit (Rad Source) with a dose rate of 1.81 Gy/min.

*Bone Marrow Chimeras*: C57BL/6NRj mice underwent total body irradiation given as two doses of 5.5 Gy irradiation, administered 6 hours apart. Following the first dose, mice received 0.1% Borgal (MSD Animal Health) in the drinking water for 2 weeks. One day after irradiation, bone marrow from *Tcf7*^DTR-GFP^ donor mice was harvested by flushing the tibiae and femora with 2.2 mL of sterile PBS. Mice were given approximately 10^6^ bone marrow cells i.v. Reconstitution was confirmed 8 weeks later by flow cytometric analysis of the blood. Chimeras were rested at least 8 weeks before use in experiments.

#### Tissue collection and processing

Tumors, tumor-draining lymph nodes (tdLNs) and spleens were excised and placed in RPMI medium supplemented with 10% FCS. Tumors were manually minced into small fragments and digested in RPMI containing 10% FCS, 1 mg/mL collagenase IV (Thermo Fisher Scientific), and 50 µg/mL DNase I (Roche) for 45 minutes at 37°C on a rotating platform. Digestion was stopped by the addition of 5 mL of ice-cold PBS. The digested tumor material was passed through a 70-μm cell strainer using the plunger of a 5-mL syringe. Cells were washed with PBS and pelleted by centrifugation at 350 *g* for 5 minutes. Red blood cells were lysed with RBC lysis buffer (17 mM Tris, pH 7.2, and 144 mM NH4Cl) for 2 minutes at room temperature. Spleens and tdLNs were passed through a 70-μm cell strainer without enzymatic digestion and washed with complete RPMI. Blood samples were collected via submental bleeding into PBS containing 2 mM EDTA.

#### Flow Cytometry

Single-cell suspensions were incubated with anti-CD16/32 antibody in PBS for 5 minutes to block Fc receptors. Surface marker staining was performed by incubating cells with 50 μL of an antibody mixture in PBS for 30 minutes at 4°C. For intracellular staining, cells were fixed and permeabilized using the Foxp3/Transcription Factor Staining Buffer Set (Invitrogen) according to the manufacturer’s instructions. Cells were incubated with the intracellular antibody mix prepared in 1X permeabilization buffer and stained in 50 μL volumes overnight at 4°C. After staining, all samples were washed once with permeabilization buffer and once with PBS before acquisition.

For experiments involving GFP as a reporter for TCF-1 expression, samples were split in two: One part was used for surface marker staining, and the other part for intracellular staining including fixation and permeabilization.

Stained samples were acquired on a Cytek Aurora 5L spectral flow cytometer for high-dimensional analysis or a CyAn ADP flow cytometer for panels of up to four colors. FCS files were analyzed using FlowJo software version 10.10.0. Data from gated populations of interest were exported as compensated FCS files and further processed in R version 4.3.2.

Preprocessed FCS files were arcSinh transformed, and raw marker intensities were quantile normalized to ensure comparability across samples. Data were visualized using Uniform Manifold Approximation and Projection (UMAP) ^65^, and unsupervised clustering was performed using RPhenograph ^66^. Data visualization and statistical analyses were conducted using the ggplot2 package in R, with Bioconductor version 3.17 used for further analysis.

#### Single-Cell RNA Sequencing

Single-cell suspensions were prepared from individual tumor samples (n = 8 per condition). Cells were labeled using the BD Single-Cell Multiplexing Kit and pooled according to their respective experimental conditions.

Intratumoral CD8^+^ T-cells were isolated as single, live, CD45^+^, CD8^+^ cells using a BD FACSAria III cell sorter. A total of 40,000 cells were loaded onto a BD Rhapsody cartridge and processed for cDNA synthesis following the BD Rhapsody Single-Cell Capture and cDNA Synthesis protocol (Doc ID: 23-22951(01)). Library amplification was performed using the BD Rhapsody System mRNA Whole Transcriptome Analysis (WTA) and Sample Tag Library Preparation Protocol (Doc ID: 23-24119(02)).

Sequencing was carried out on an Illumina NovaSeq 6000 or NovaSeq X Plus platform using 150 bp paired-end reads (R1 = 51 cycles, R2 = 71 cycles), with an average sequencing depth of approximately 50,000 reads per cell. Data processing, including read alignment, cell barcode demultiplexing, read deduplication, expression matrix

generation, and quality reporting, was performed using the Rhapsody Sequence Analysis Pipeline v2.0.

Further downstream analysis of expression matrices was performed in R v4.3.2. Cells were filtered to exclude those with more than 10% mitochondrial content or less than 2% ribosomal protein content. Duplicate cells were identified and removed using scDblFinder ^67^. The Seurat pipeline v5.1 ^68^ was used for normalization, dimensionality reduction, and clustering. CD8^+^ T-cell populations and their subsets were annotated using ProjectTILs ^33^ and scGate ^69^. Gene set enrichment analysis for T-cell programs was performed with the UCell package ^34^.

To investigate cellular trajectories and estimate pseudotime, Monocle 3 ^36,37^ was used. Cells with the highest expression of *Sell* and *Tcf7* were chosen as the starting point for trajectory inference. Monocle 3 then arranged intratumoral CD8^+^ T-cells along a trajectory according to their transcriptional progression, with default settings applied throughout the analysis.

#### Statistical analysis

One-way ANOVA followed by Tukey’s post-hoc test was used to compare three or more groups. For tumor size comparisons, the last time point was used for the one-way ANOVA analysis. For comparisons between two groups, the Mann-Whitney U-test was used. A *p*- value < 0.05 was considered significant. All statistical analyses were performed using GraphPad Prism software (v10.0.0).

## Key Resources Table

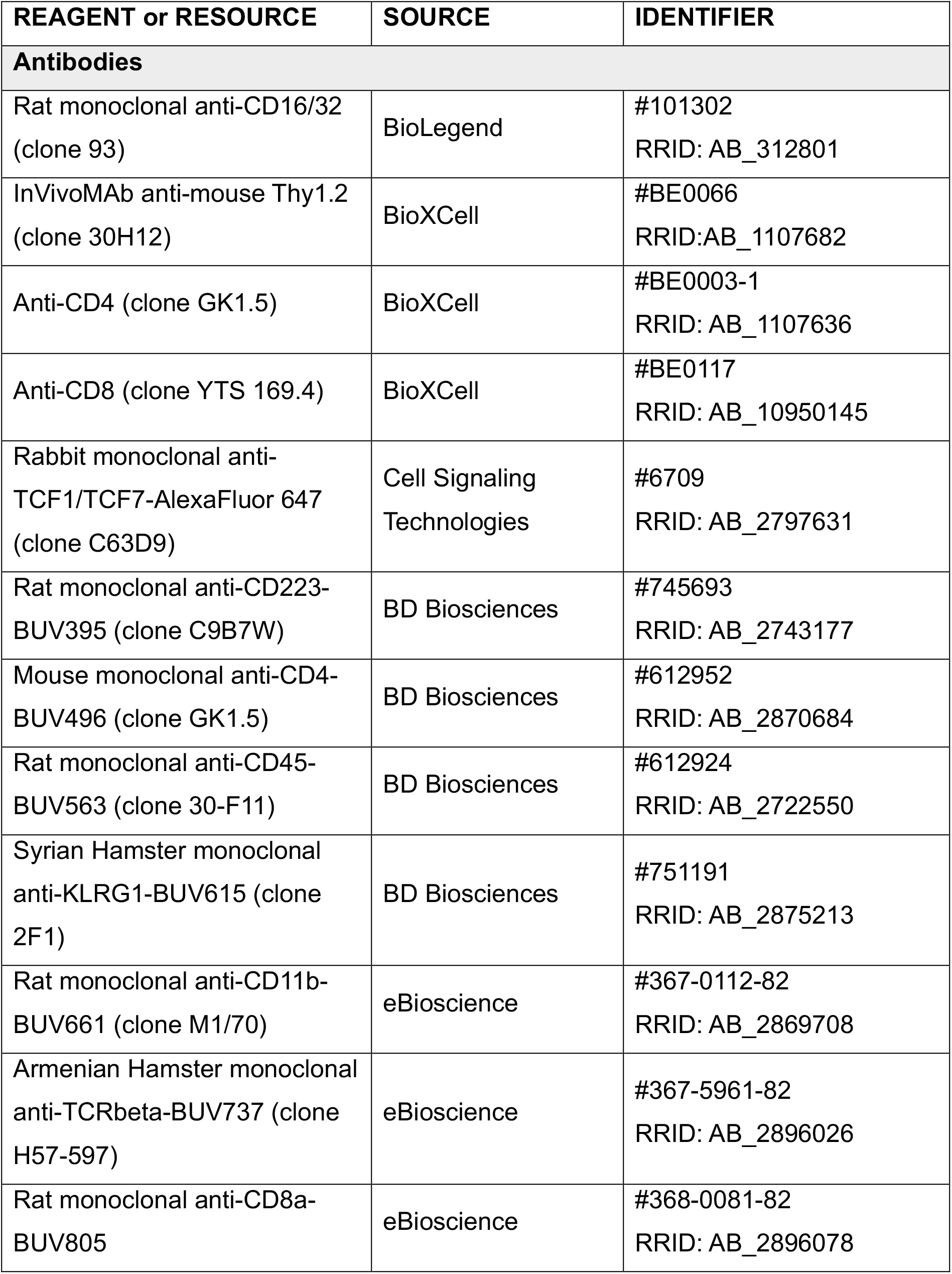
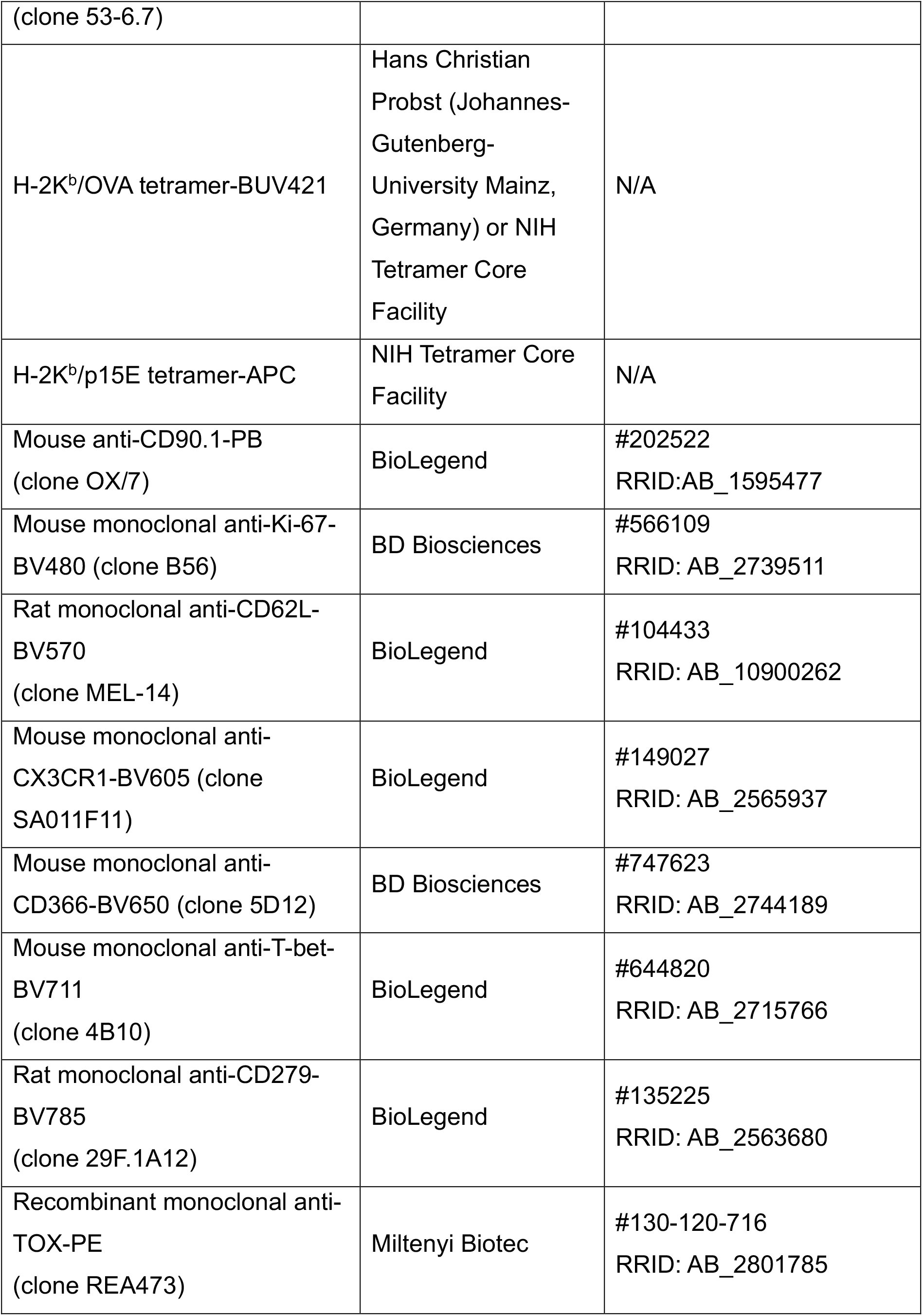
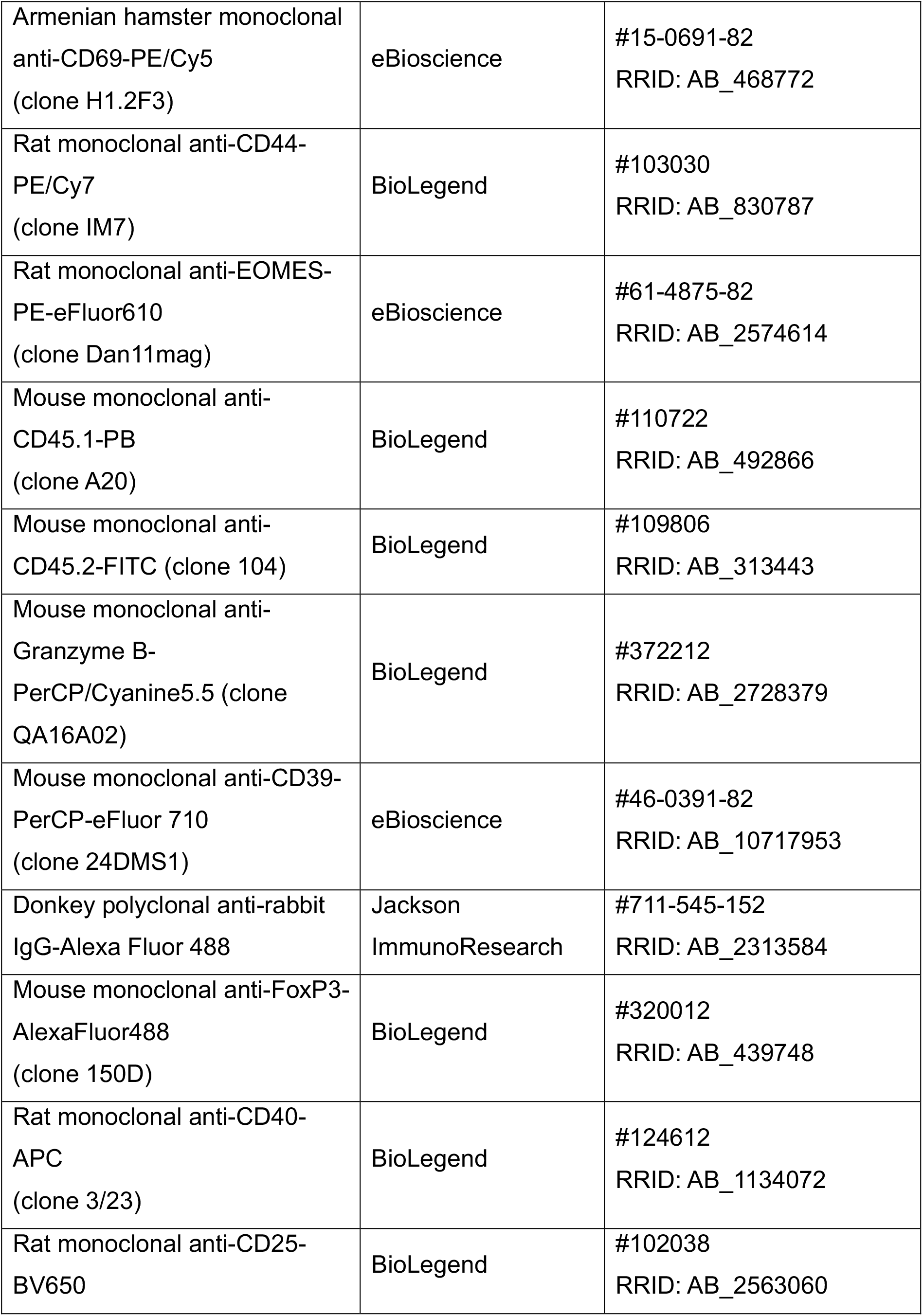
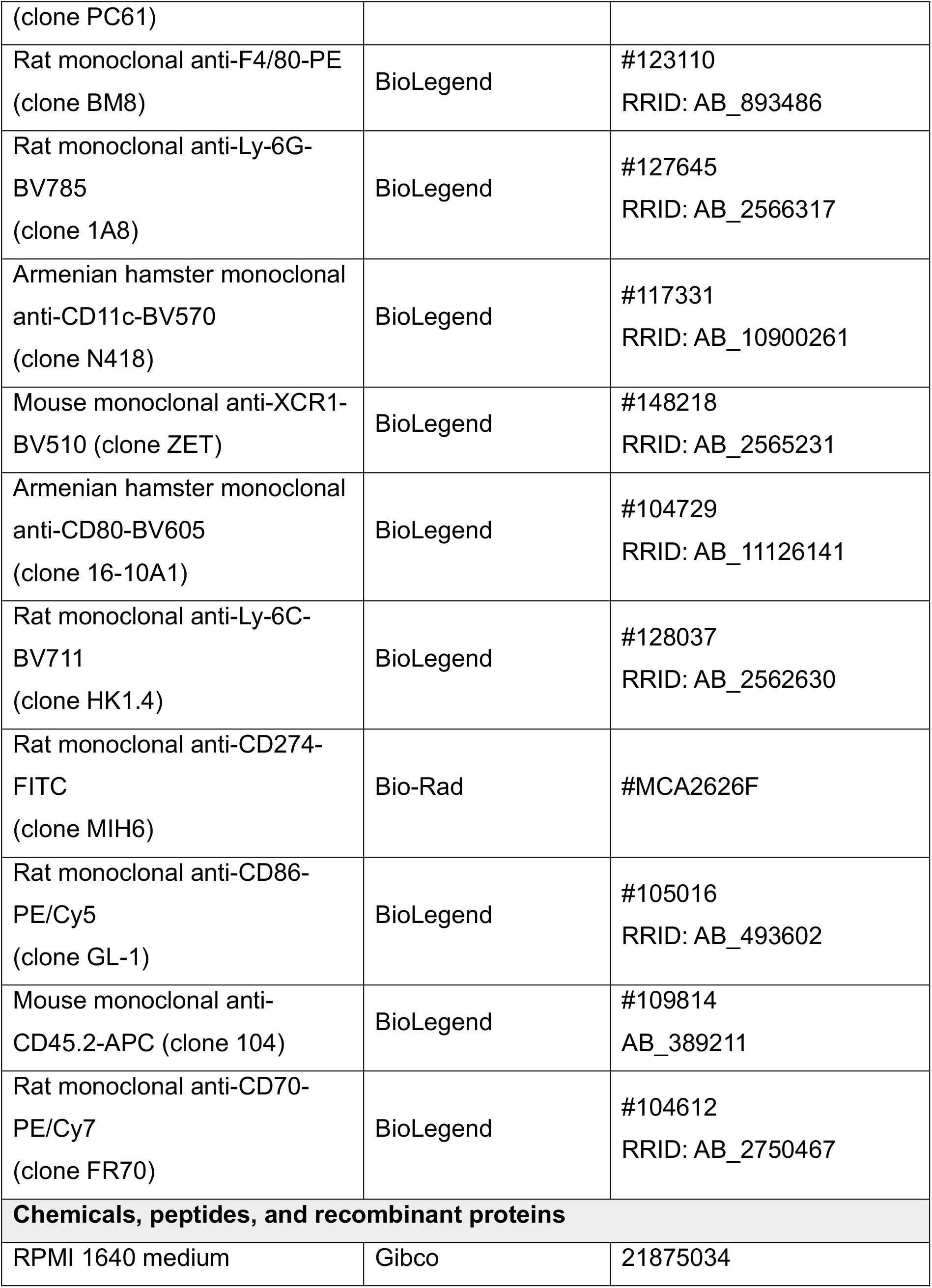
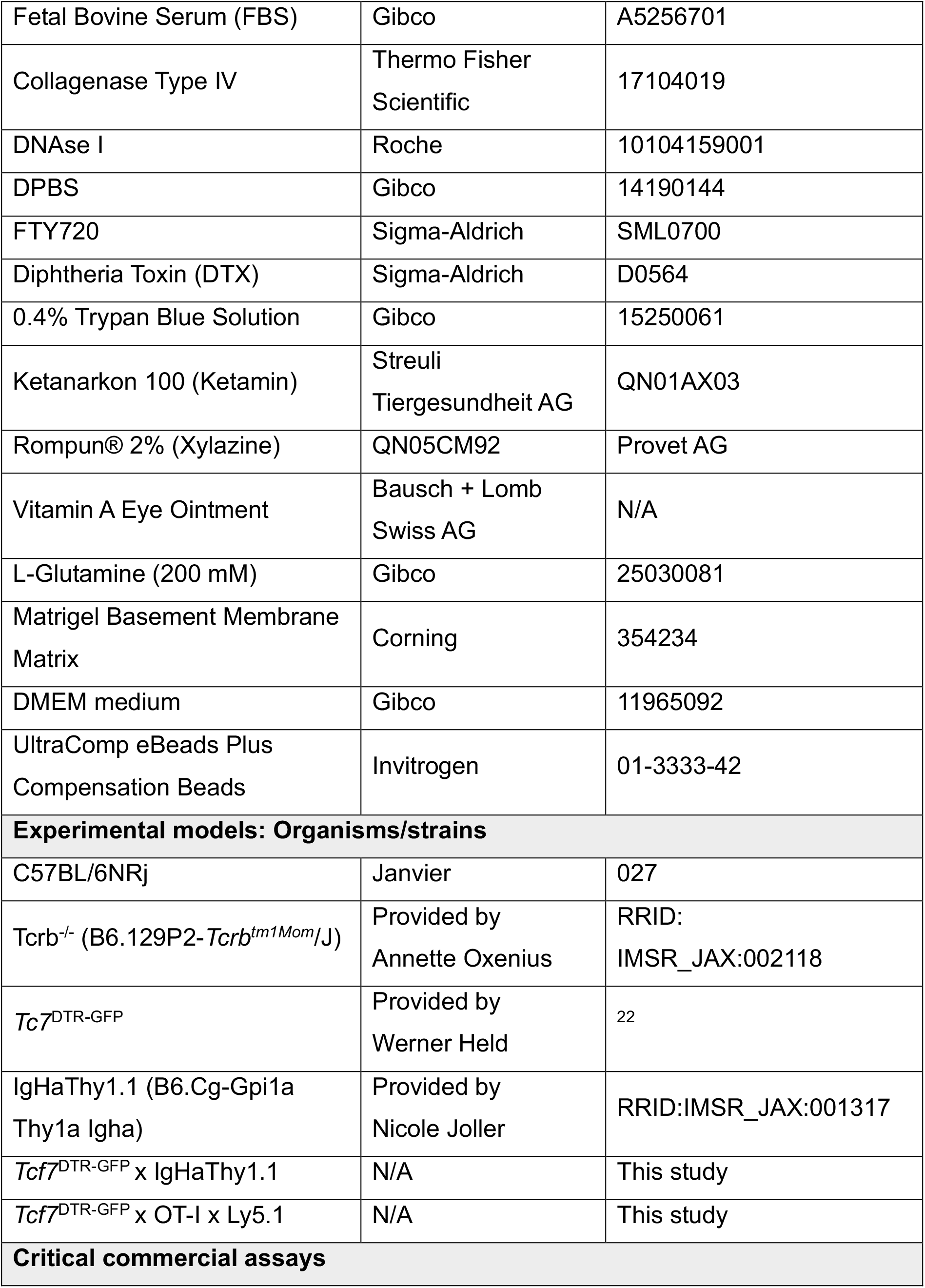
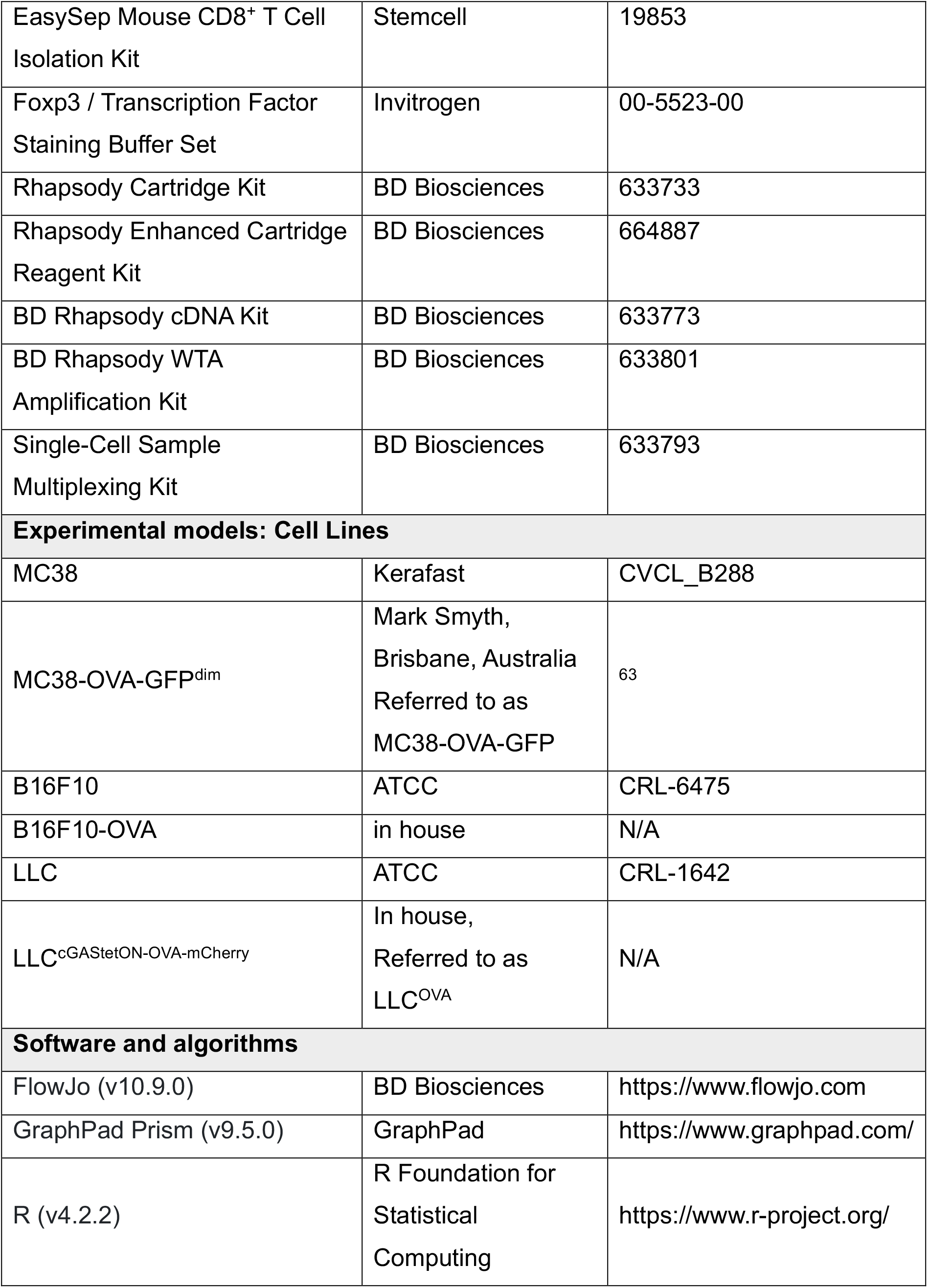
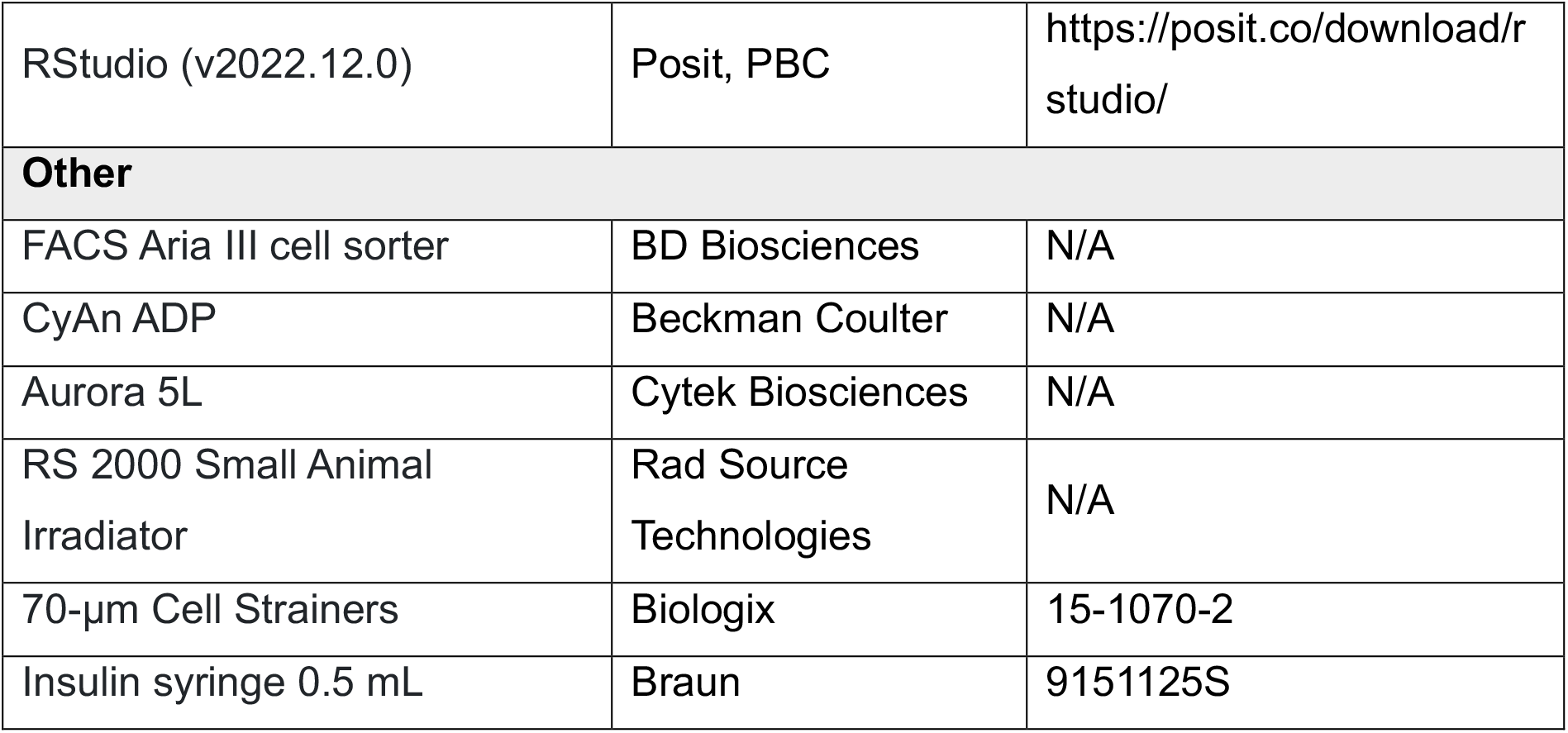

---

**Figure S1.**
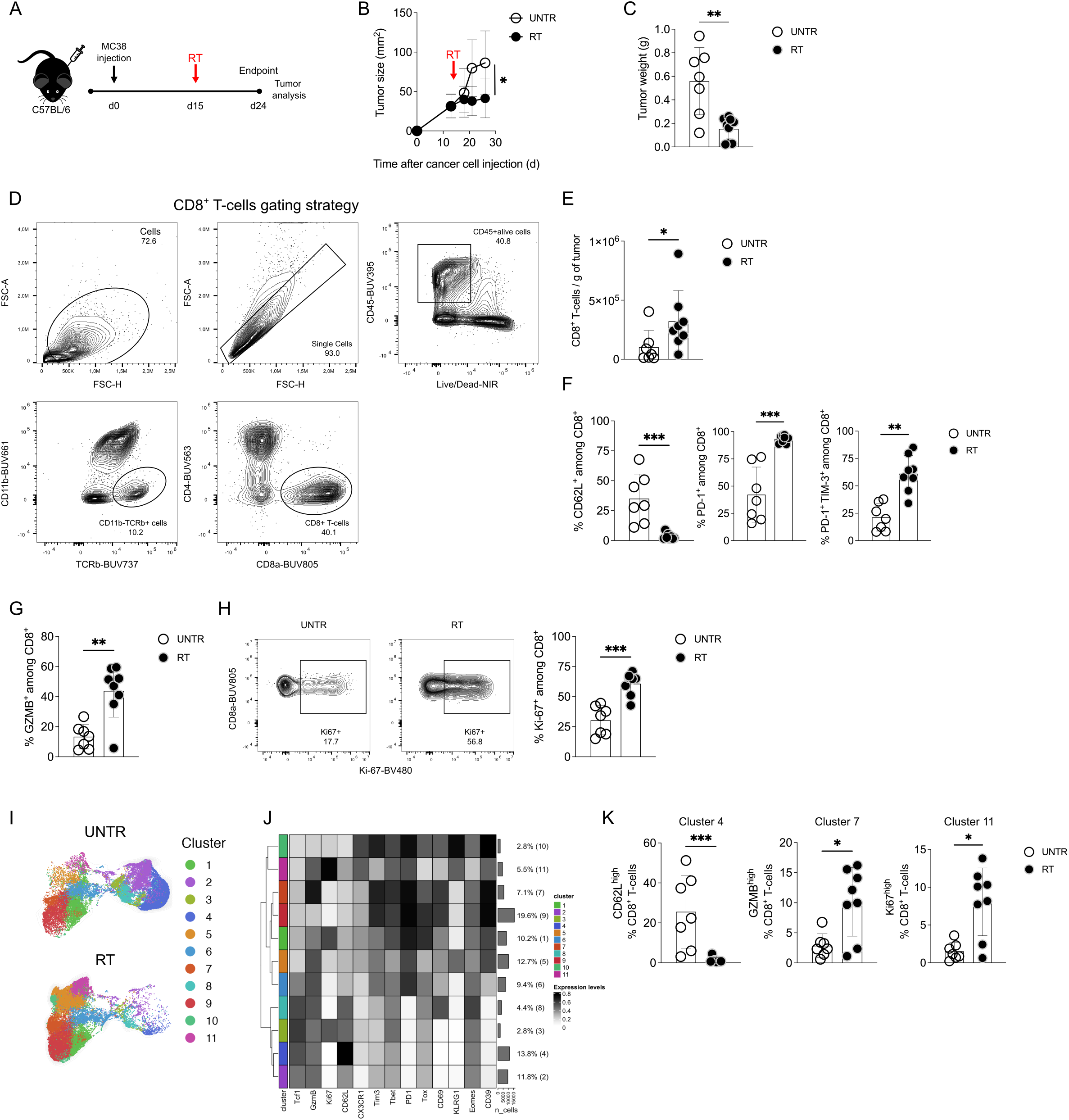
Radiotherapy induces differentiation of intratumoral CD8+ T-cells. (A) Experimental setup: C57BL/6 mice (UNTR n=7, RT n=9) were injected subcutaneously with 6x105 MC38 cells in matrigel:PBS (1:2). Fifteen days later, some groups received radiotherapy (RT; 20 Gy) or not (UNTR). Tumors were analyzed nine days after radiotherapy. (B) Tumor size (length x breadth) was measured by caliper. (C) Tumor weight. (D) Flow cytometry gating strategy for intratumoral CD8+ T-cells. (E) Number of CD8+ T-cells per gram of tumor. (F) Percentage of CD62L, PD-1, PD-1 and TIM-3 expressing CD8+ T-cells. (G) Percentage of GZMB-expressing CD8+ T-cells. (H) Percentage of Ki-67 expressing CD8+ T-cells. (I) Uniform manifold approximation and Projection (UMAP) representation of CD8+ T-cell clusters. (J) The heatmap illustrates the relative expression of key markers across different clusters. (K) Percentage of UMAP clusters 4, 7 and 11. Each symbol represents one mouse. The bar indicates the mean ± SD. Groups were statistically compared using the Mann-Whitney U test. *p<0.05, **p<0.01, ***p<0.001.

**Figure S2.**
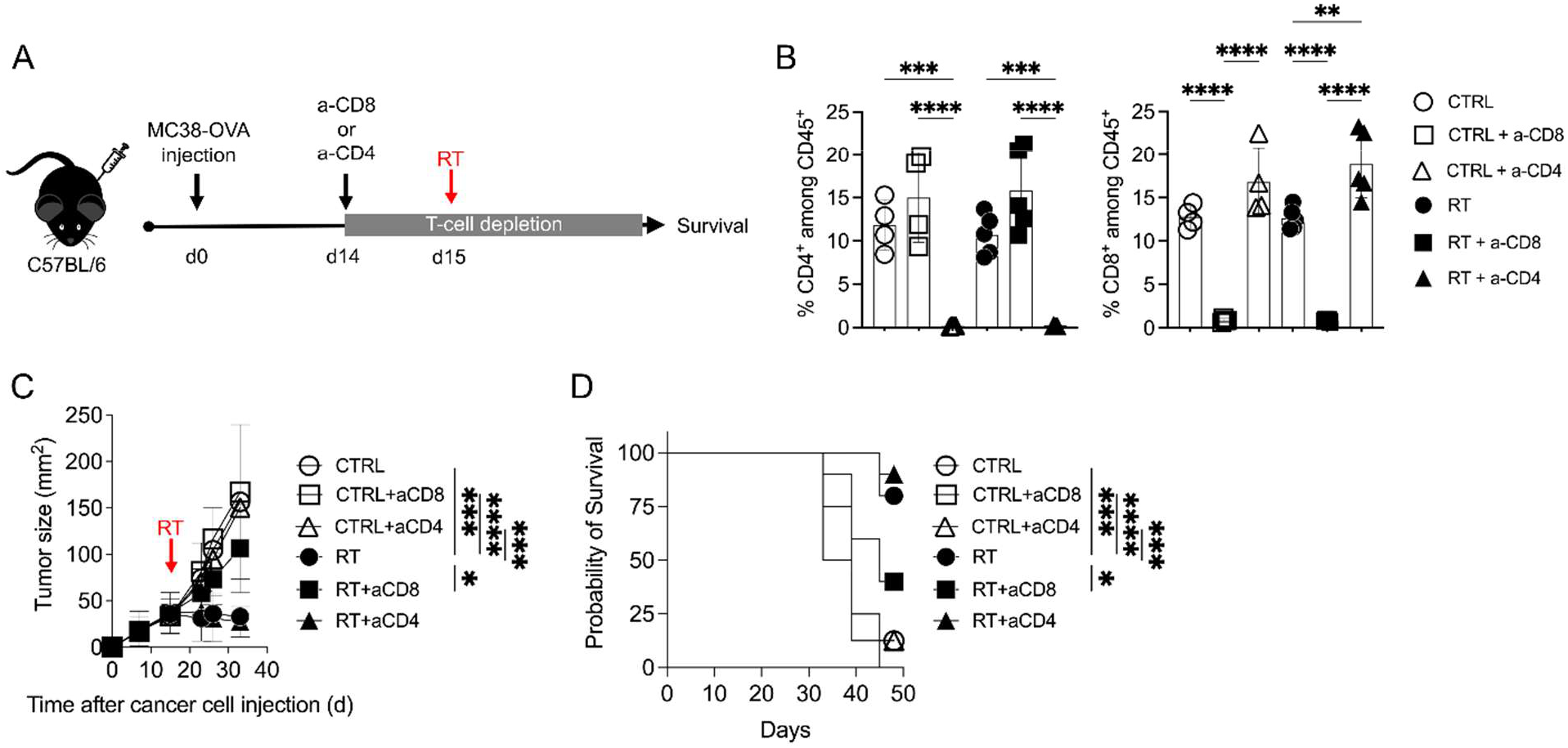
The therapeutic efficacy of radiotherapy depends on CD8+ T-cells. (A) Experimental setup: C57BL/6 mice (UNTR n=8 per group, RT n=10 per group) were injected subcutaneously with 8x105 MC38-OVA cells in matrigel:PBS (1:2). Fifteen days later, some groups received radiotherapy (RT; 20 Gy) or not (UNTR). Starting 1 day before radiotherapy, CD4 or CD8 cells were depleted by intraperitoneal injection of 500 µg of anti-CD4 (GK1.5) or anti-CD8 (YTS169.4), followed by a second dose 12 days after radiotherapy. (B) Depletion validation was performed two days after antibody injection by measuring percentages of CD4+ and CD8+ cells among CD45+ cells in the blood. (C) Tumor size (length x breadth) was measured by caliper. Each symbol represents one mouse. The bar indicates the mean ± SD. Groups were statistically compared using one-way ANOVA with Tukey’s multiple comparison correction. (D) Kaplan-Meier survival curves. Groups were statistically compared using the Mantel-Cox test. *p<0.05, **p<0.01, ***p<0.001, ****p<0.0001.

**Figure S3.**
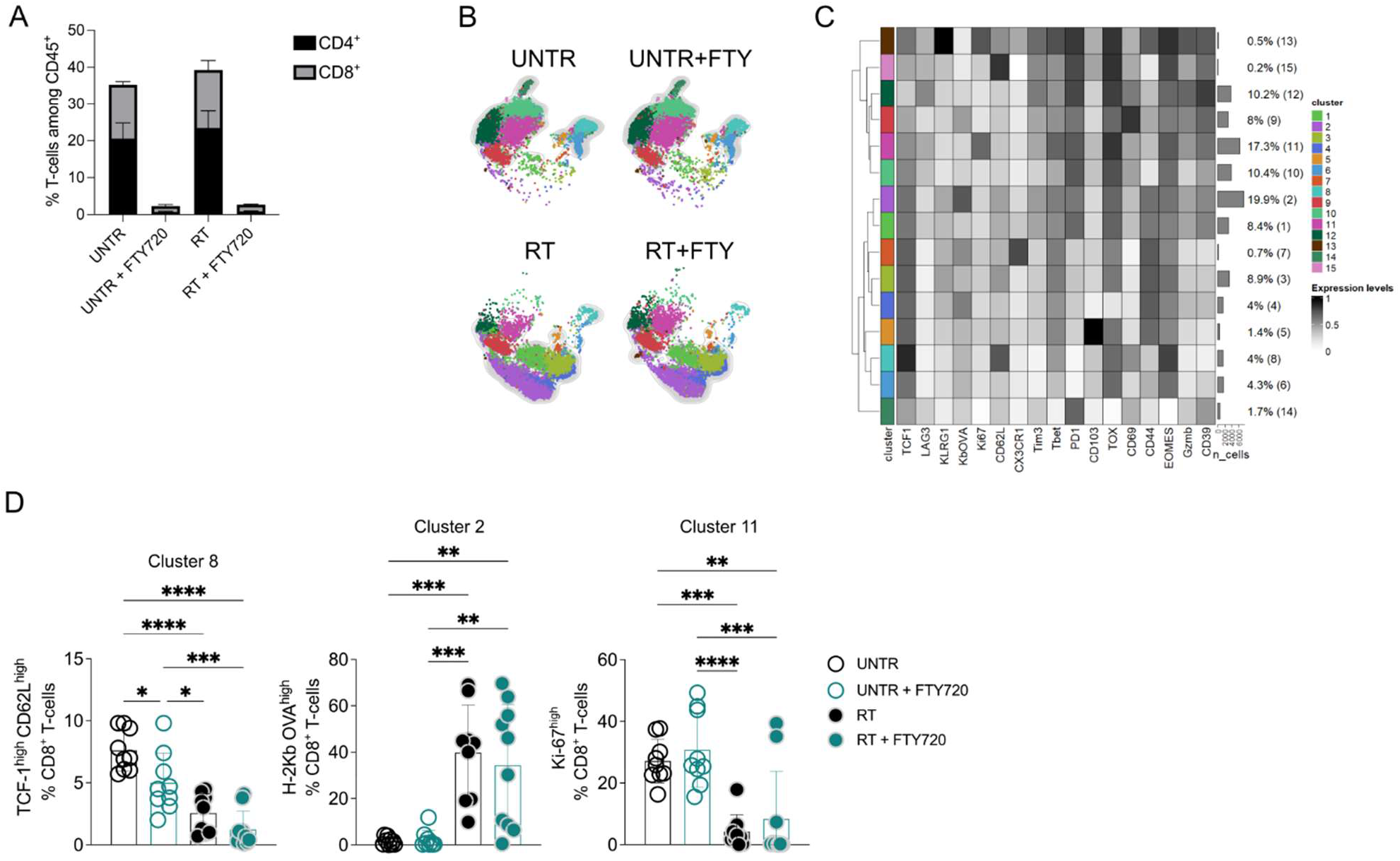
Radiotherapy increases the frequency of tumor-responsive intratumoral CD8+ T-cell clusters. (A) FTY720 validation was performed two days after radiotherapy by measuring percentages of CD4+ and CD8+ cells among CD45+ cells in the blood. (B) UMAP representation of CD8+ T-cell clusters. (C) The heatmap illustrates the relative expression of key markers across different clusters. (D) Percentage of UMAP clusters 8, 2 and 11. The bar indicates the mean ± SD. Groups were statistically compared using one-way ANOVA with Tukey’s multiple comparison correction. *p<0.05, **p<0.01, ***p<0.001, ****p<0.0001.

**Figure S4.**
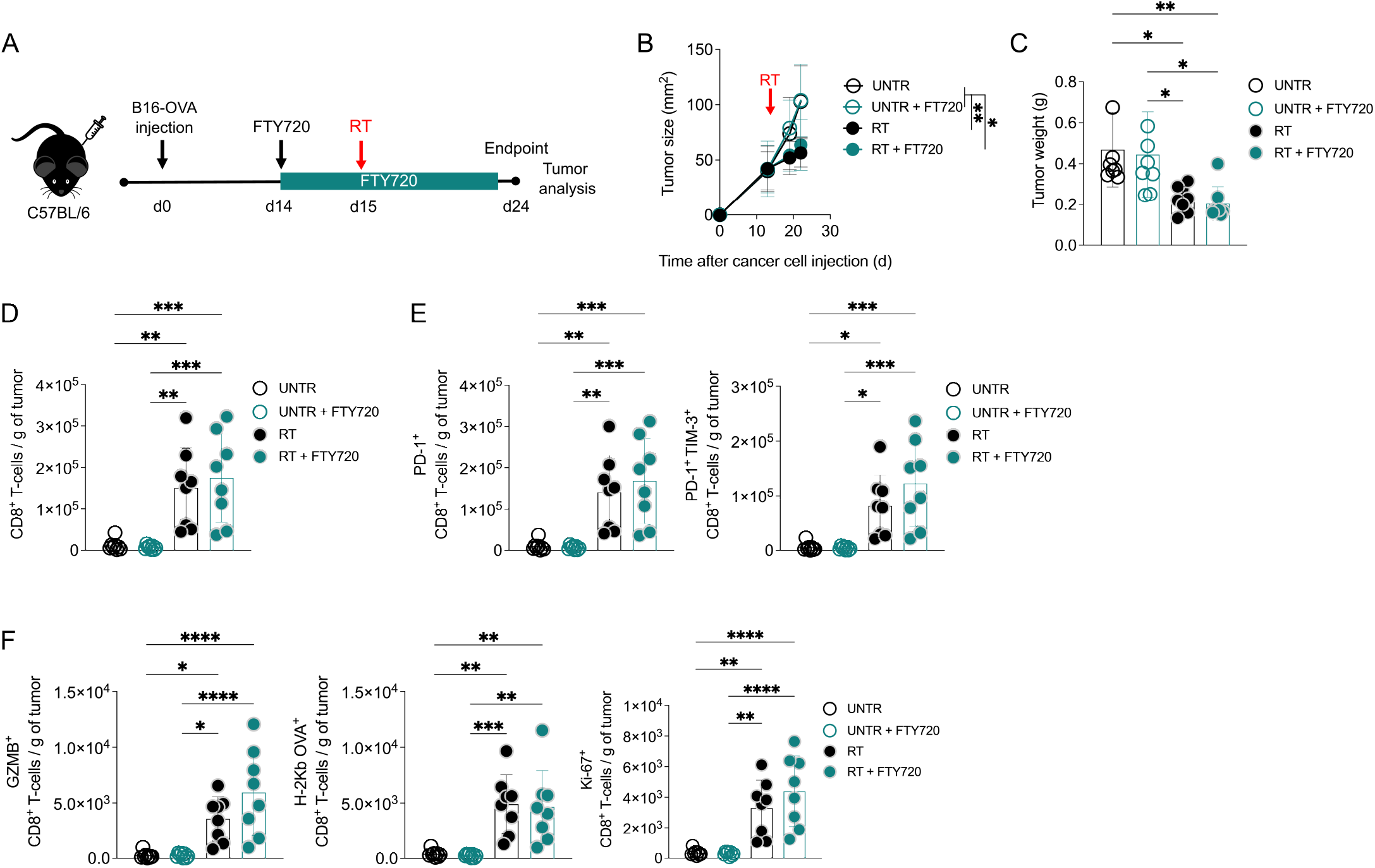
Pre-existing intratumoral CD8+ T-cells are sufficient for the therapeutic efficacy of radiotherapy. (A) Experimental setup: C57BL/6 mice (n=8 per condition) were injected subcutaneously with 8x105 MC38-OVA cells in matrigel:PBS (1:2). Fifteen days later, some groups received radiotherapy (RT; 20 Gy) or not (UNTR). Starting 1 day before radiotherapy, some groups received 20 µg FTY720 per os every second day until the endpoint. Tumors were analyzed nine days after radiotherapy. (B) Tumor size (length x breadth) was measured by caliper. (C) Tumor weight (D) Number of CD8+ T- cells per gram of tumor. (E) Number of PD-1, PD-1 and TIM-3 expressing CD8+ T-cells per gram of tumor. (F) Number of GZMB and Ki-67 expressing CD8+ T-cells and SIINFEKL-specific (determined by H-2Kb OVA tetramer) CD8+ T-cells per gram of tumor, determined by H-2Kb OVA tetramer. The bar indicates the mean ± SD. Groups were statistically compared using one-way ANOVA with Tukey’s multiple comparison correction. *p<0.05, **p<0.01, ***p<0.001, ****p<0.0001.

**Figure S5.**
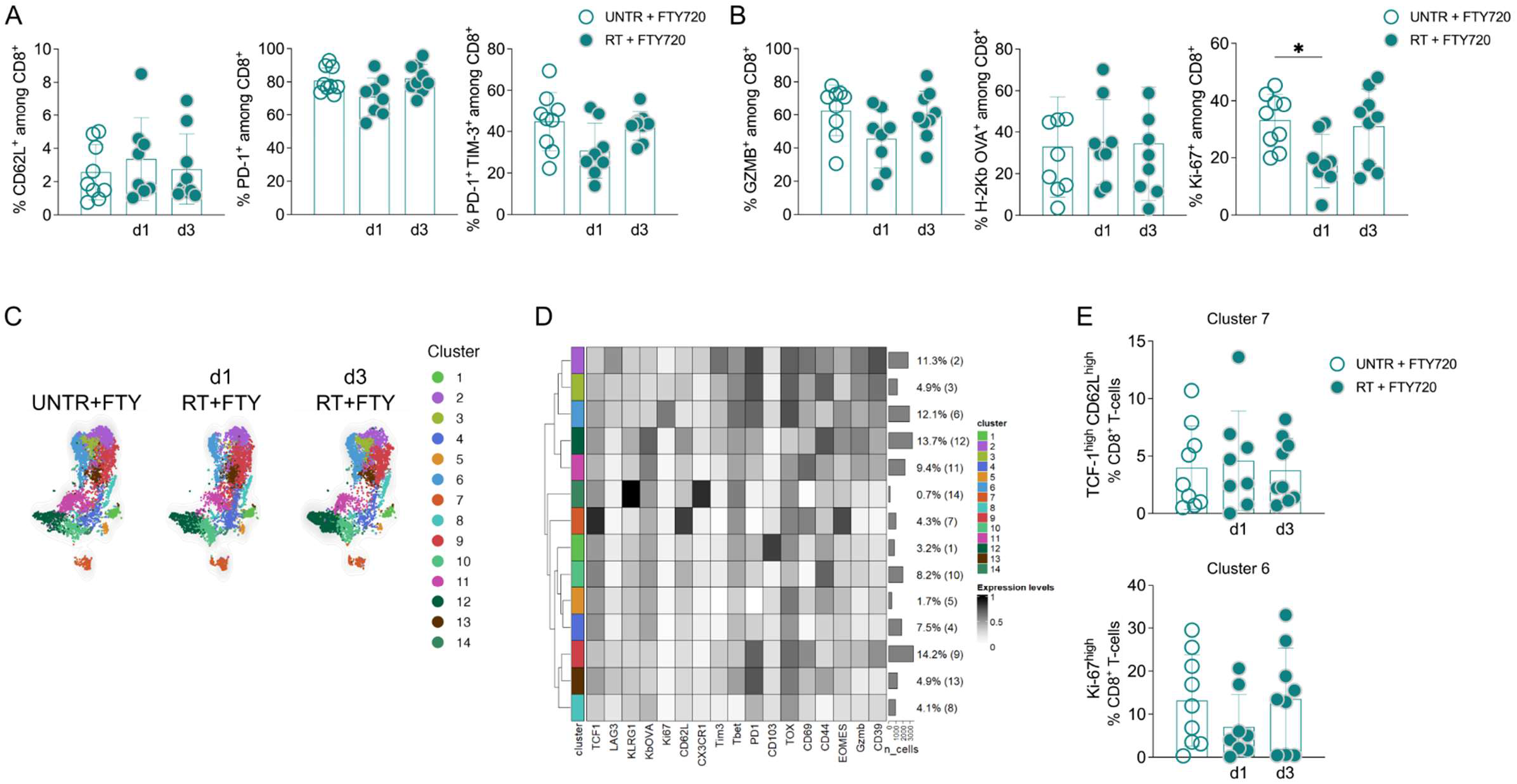
Radiotherapy initially decreases the frequency of proliferating intratumoral CD8+ T-cells. (A) Percentage of CD62L, PD-1, PD-1 and TIM-3 expressing CD8+ T-cells. (B) Percentage of GZMB and Ki-67 expressing CD8+ T-cells and SIINFEKL-specific CD8+ T-cells, determined by H-2Kb OVA tetramer. (C) UMAP representation of CD8+ T-cell clusters. (D) The heatmap illustrates the relative expression of key markers across different clusters. (E) Percentage of UMAP clusters 6 and 7. The bar indicates the mean ± SD. Groups were statistically compared using one-way ANOVA with Tukey’s multiple comparison correction. *p<0.05.

**Figure S6.**
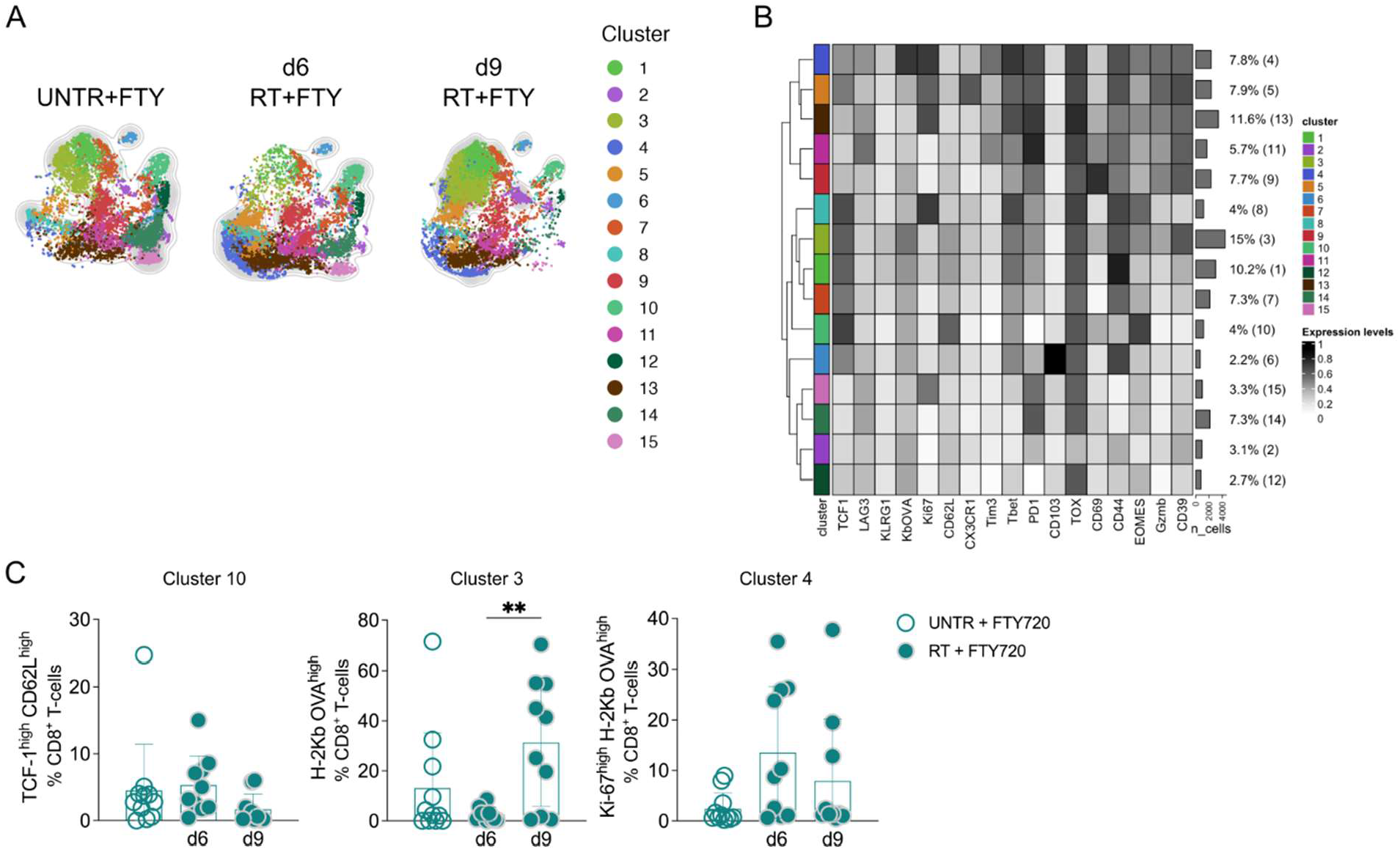
Radiotherapy leads to an increased frequency of proliferating intratumoral CD8+ T-cell clusters. (A) UMAP representation of CD8+ T-cell clusters. (B) The heatmap illustrates the relative expression of key markers across different clusters. (C) Percentage of UMAP clusters 10, 3 and 4. The bar indicates the mean ± SD. Groups were statistically compared using one-way ANOVA with Tukey’s multiple comparison correction. **p<0.01.

**Figure S7.**
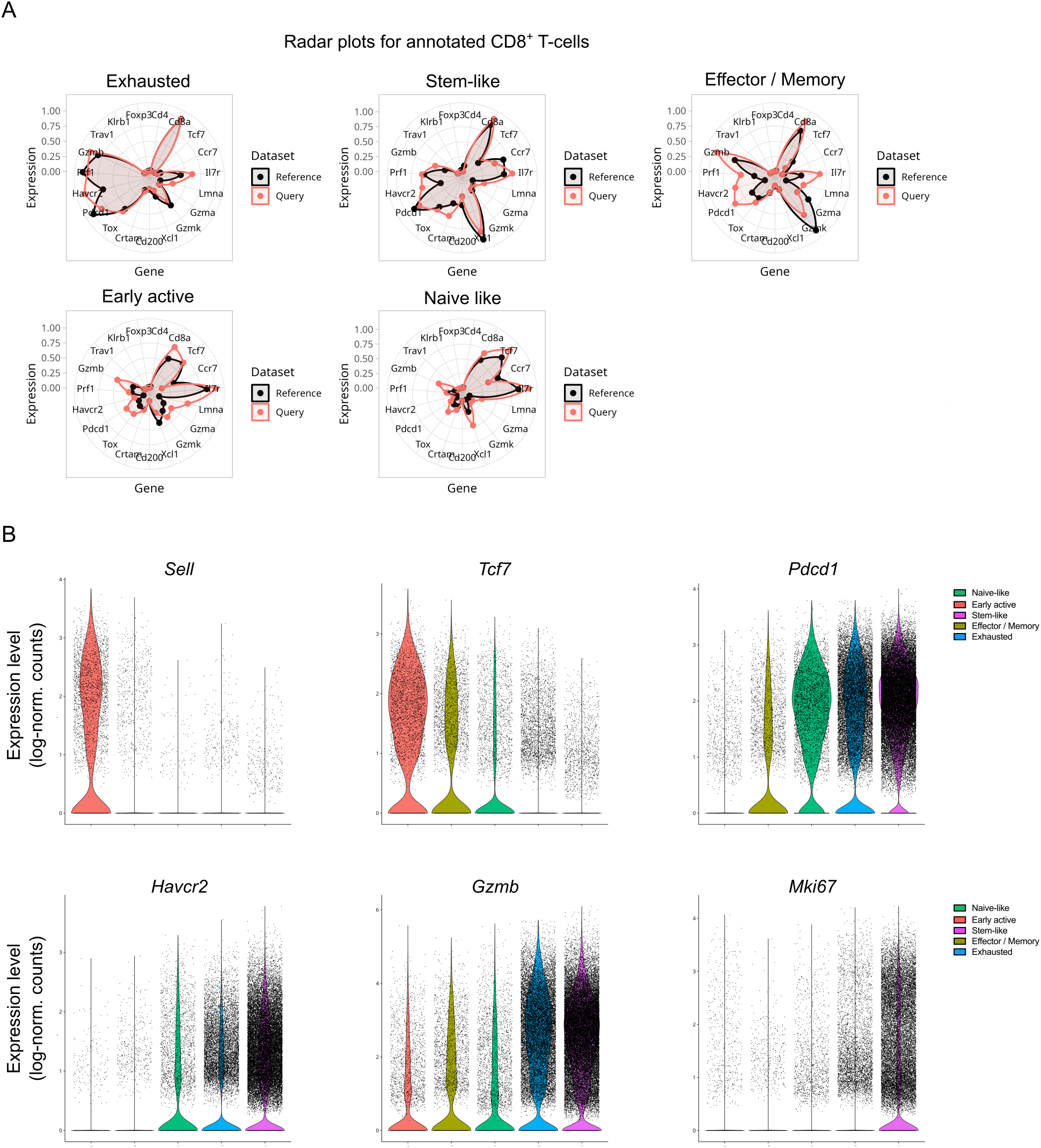
Validation of ProjectTILs CD8+ T-cell annotation. (A) Radar plots showing the expression profiles of key differentiation and activation markers for CD8+ T- cells. Reference samples (black) and pooled experimental samples (red) are overlaid for each subset. (B) Expression of key differentiation and activation markers presented as log-normalized counts for each CD8+ T-cell subset.

**Figure S8.**
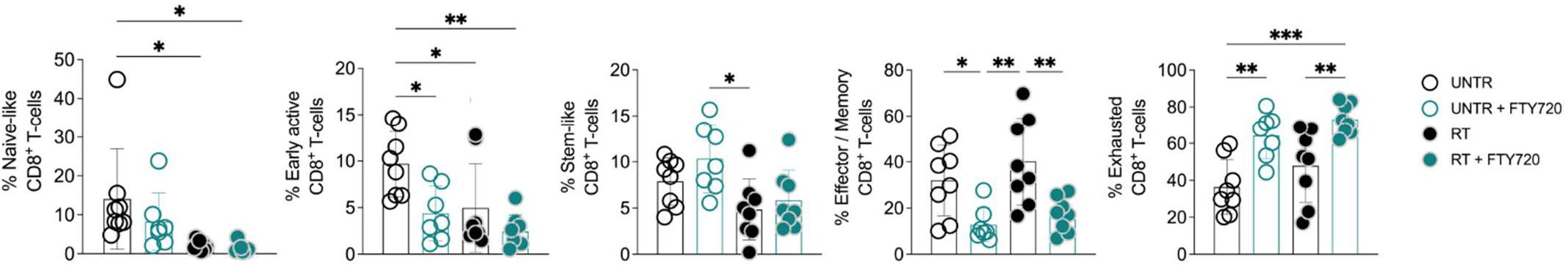
Radiotherapy leads to an increased frequency of pre-existing exhausted intratumoral CD8+ T-cells. Percentage of ProjectTILs annotated naïve-like, early active, stem-like, effector/memory and exhausted intratumoral CD8+ T-cells. The bar indicates the mean ± SD. Groups were statistically compared using one-way ANOVA with Tukey’s multiple comparison correction. *p<0.05, **p<0.01, ***p<0.001.

**Figure S9.**
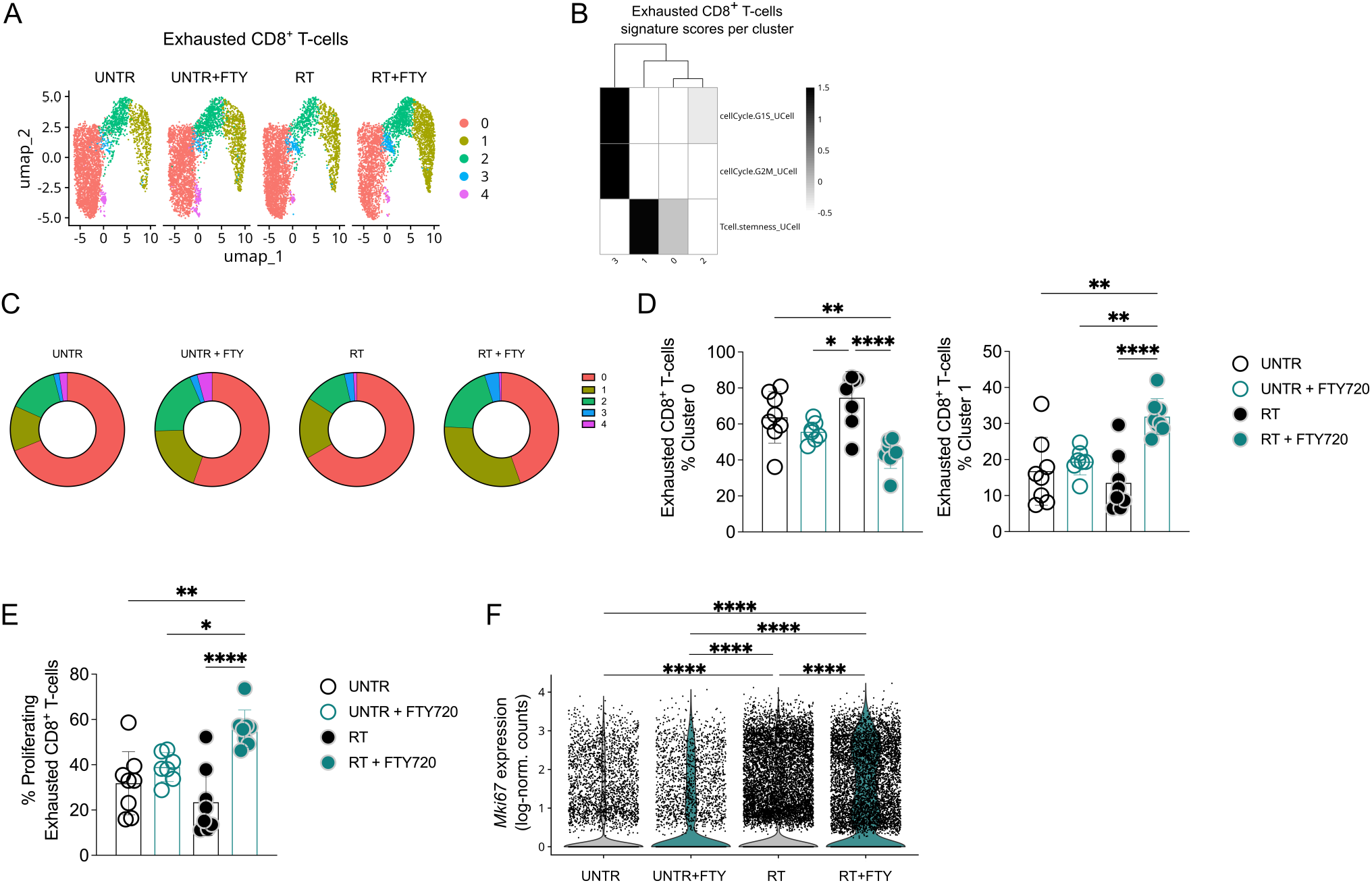
Radiotherapy promotes the proliferation of pre-existing intratumoral exhausted CD8+ T-cells. (A) UMAP representation of exhausted CD8+ T-cell clusters. (B) Heatmap illustrates signature scores (determined by UCell package) across different exhausted CD8+ T-cell clusters. (C) Pie chart showing frequencies of exhausted CD8+ T-cell subsets per condition. (D) Percentage of UMAP clusters 0 and 1. (E) Percentage of proliferating exhausted CD8+ T-cells (defined by a G1S or G2M score >0.1), determined by UCell package. (F) Mki67 expression shown as log-normalized counts. The bar indicates the mean ± SD. Groups were statistically compared using one-way ANOVA with Tukey’s multiple comparison correction. *p<0.05, **p<0.01, ****p<0.0001.

**Figure S10.**
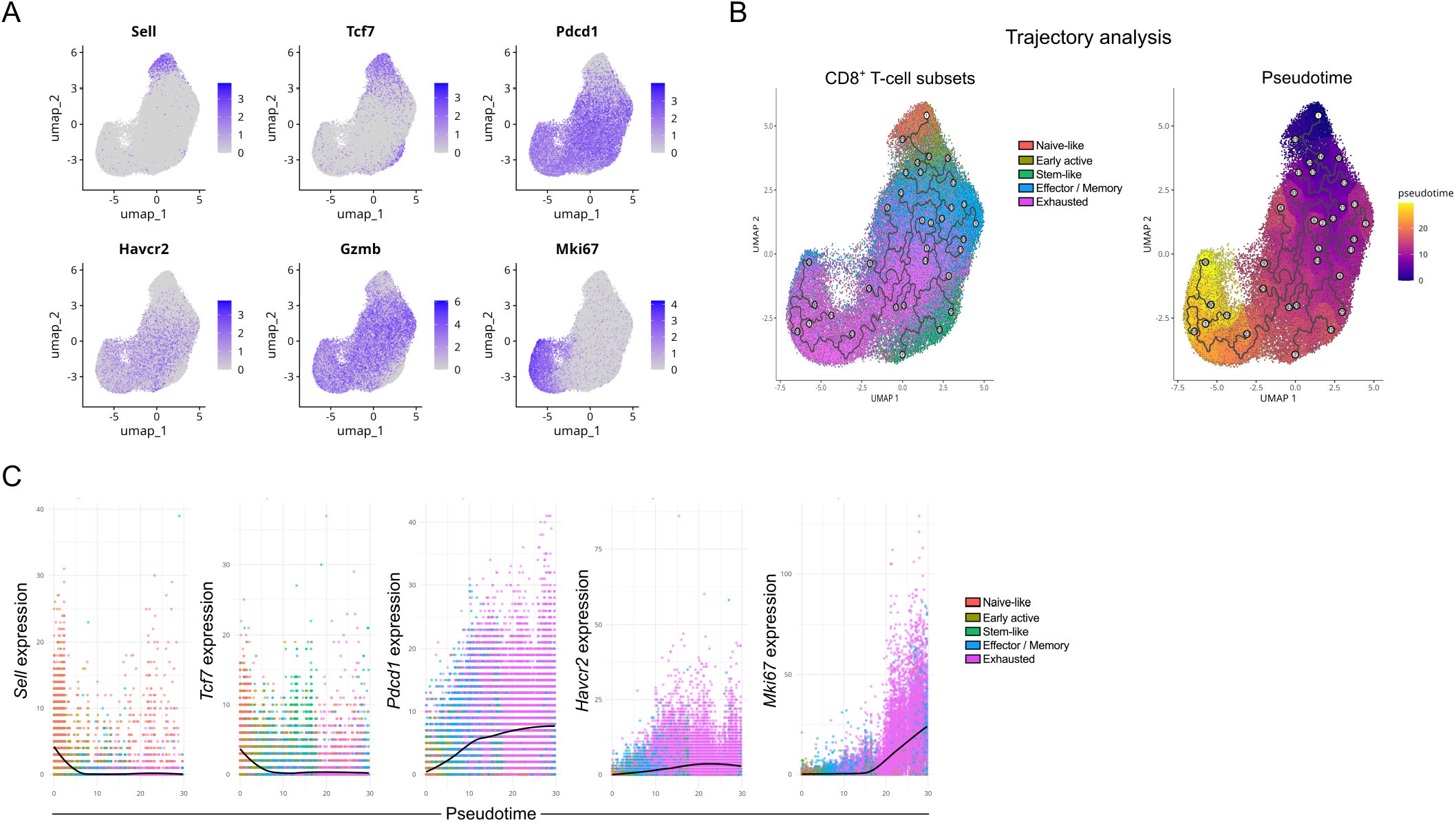
Trajectory analysis identifies intratumoral stem-like CD8+ T-cells as a potential reserve of exhausted CD8+ T-cells. (A) UMAP plots showing the expression of key activation and differentiation genes across CD8+ T-cells. (B) Pseudotime trajectory of CD8+ T-cells visualized with Monocle3. Cells are colored by CD8+ T-cell subsets (left) or pseudotime (right), with the trajectory graph displayed, excluding branch points. (C) Scatter plots display the expression of key CD8+ T-cell differentiation and activation genes across pseudotime, with cells colored by CD8+ T-cell subsets. Each plot includes a trendline to illustrate the general expression trajectory.

**Figure S11.**
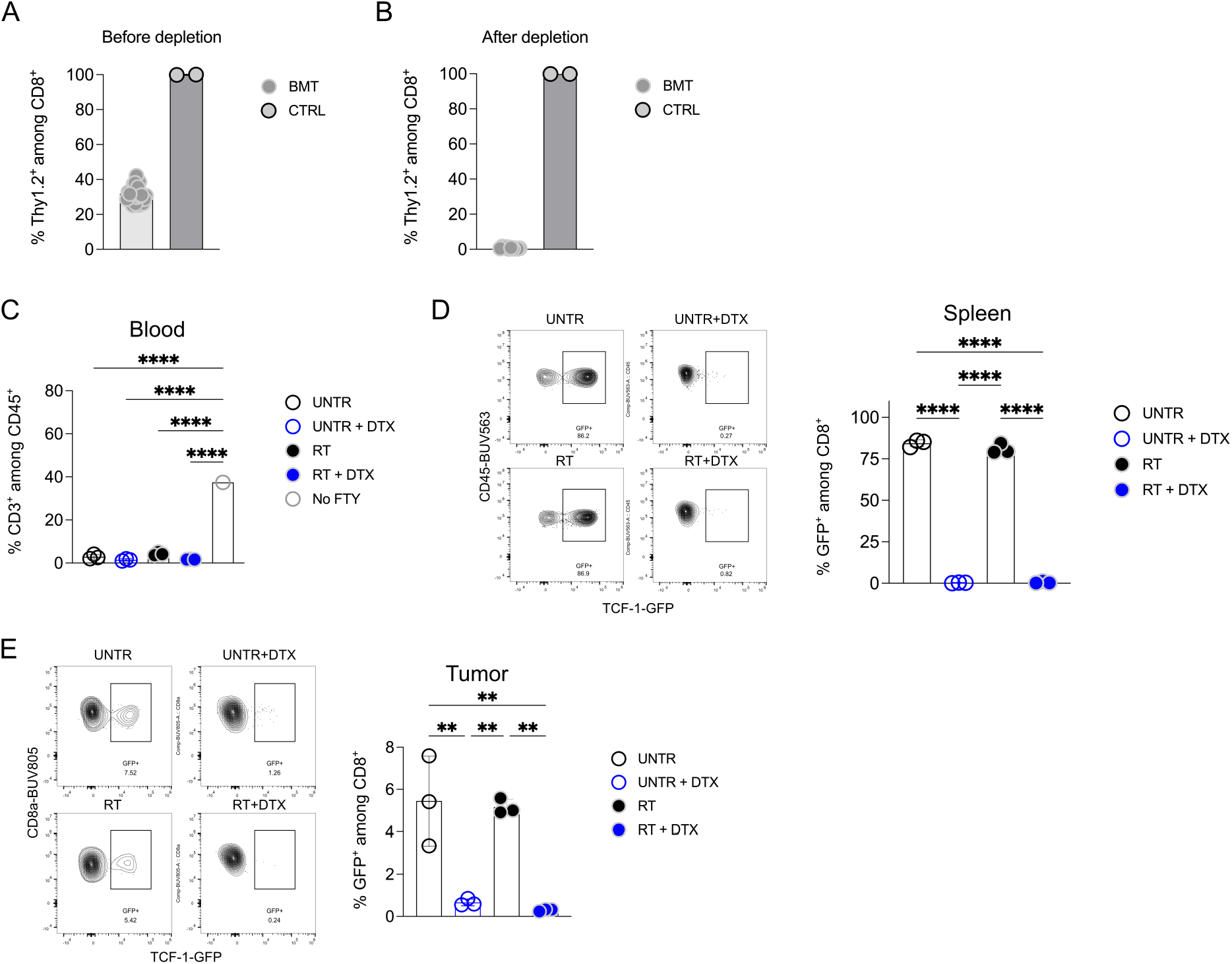
Validation of TCF-1+ depletion in spleen and tumor. (A-B) Host T-cell (Thy1.2+) depletion was validated by measuring the percentage of Thy1.2+ cells among CD8+ cells in the blood two days before (A) and after (B) the first injection. (C) FTY720 validation was performed by measuring the percentage of CD3+ cells among CD45+ cells in the blood one day after radiotherapy. (D-E) Depletion of TCF-1+ (GFP+) cells was validated by measuring the percentage of GFP+ cells among CD8+ cells in the spleen (D) and unfixed tumor samples (E) at the endpoint. The bar indicates the mean ± SD. Each symbol represents one mouse. Groups were statistically compared using one-way ANOVA with Tukey’s multiple comparison correction. **p<0.01, ****p<0.0001.

**Figure S12.**
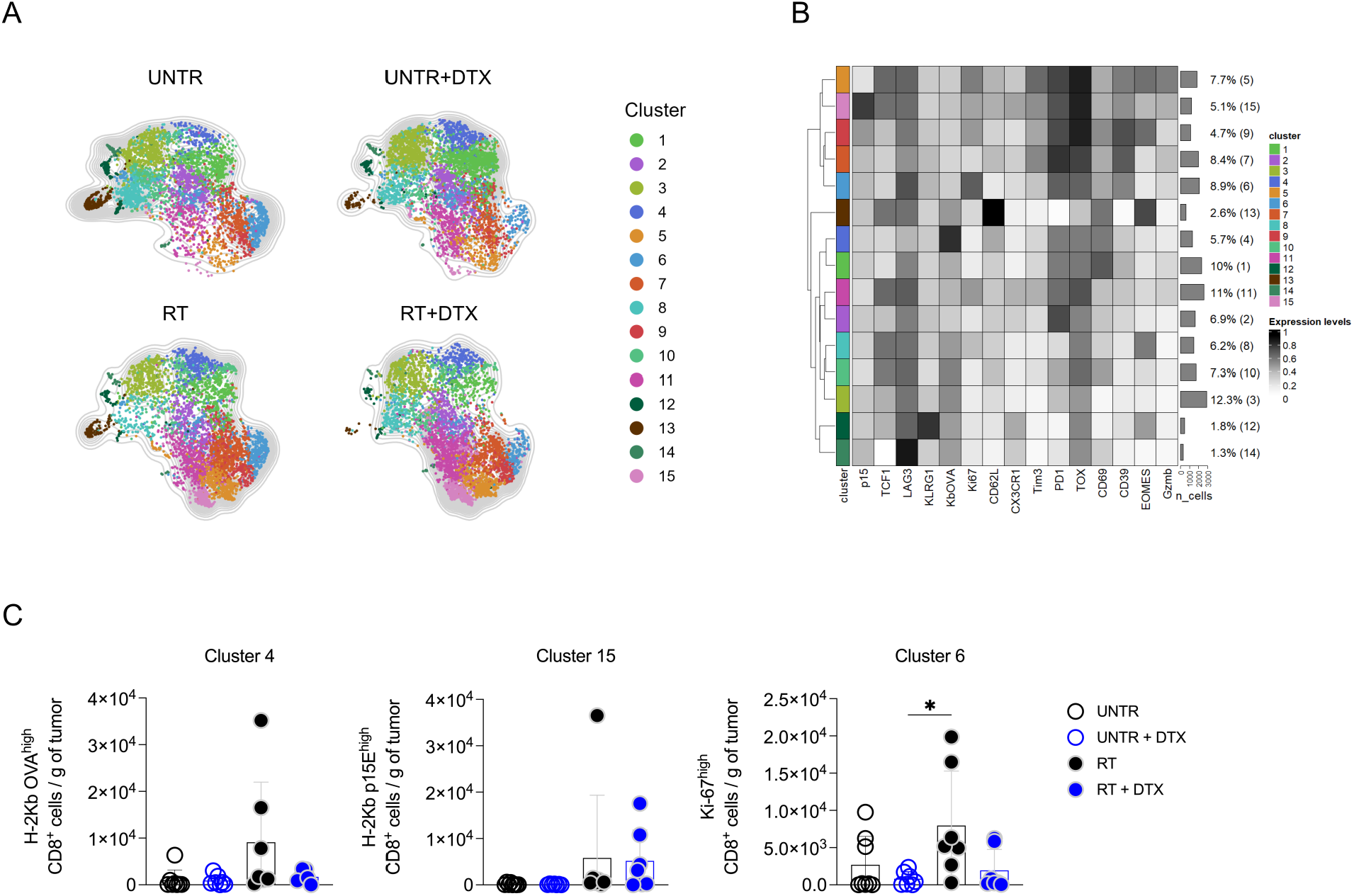
TCF-1+ cell depletion reduces counts of tumor-responsive CD8+ T-cell clusters after radiotherapy. (A) UMAP representation of CD8+ T-cell clusters. (B) The heatmap illustrates the relative expression of key markers across different clusters. (C) Number of UMAP clusters 4, 15 and 6 per gram of tumor. Each symbol represents one mouse. Groups were statistically compared using one-way ANOVA with Tukey’s multiple comparison correction. *p<0.05.

**Figure S13.**
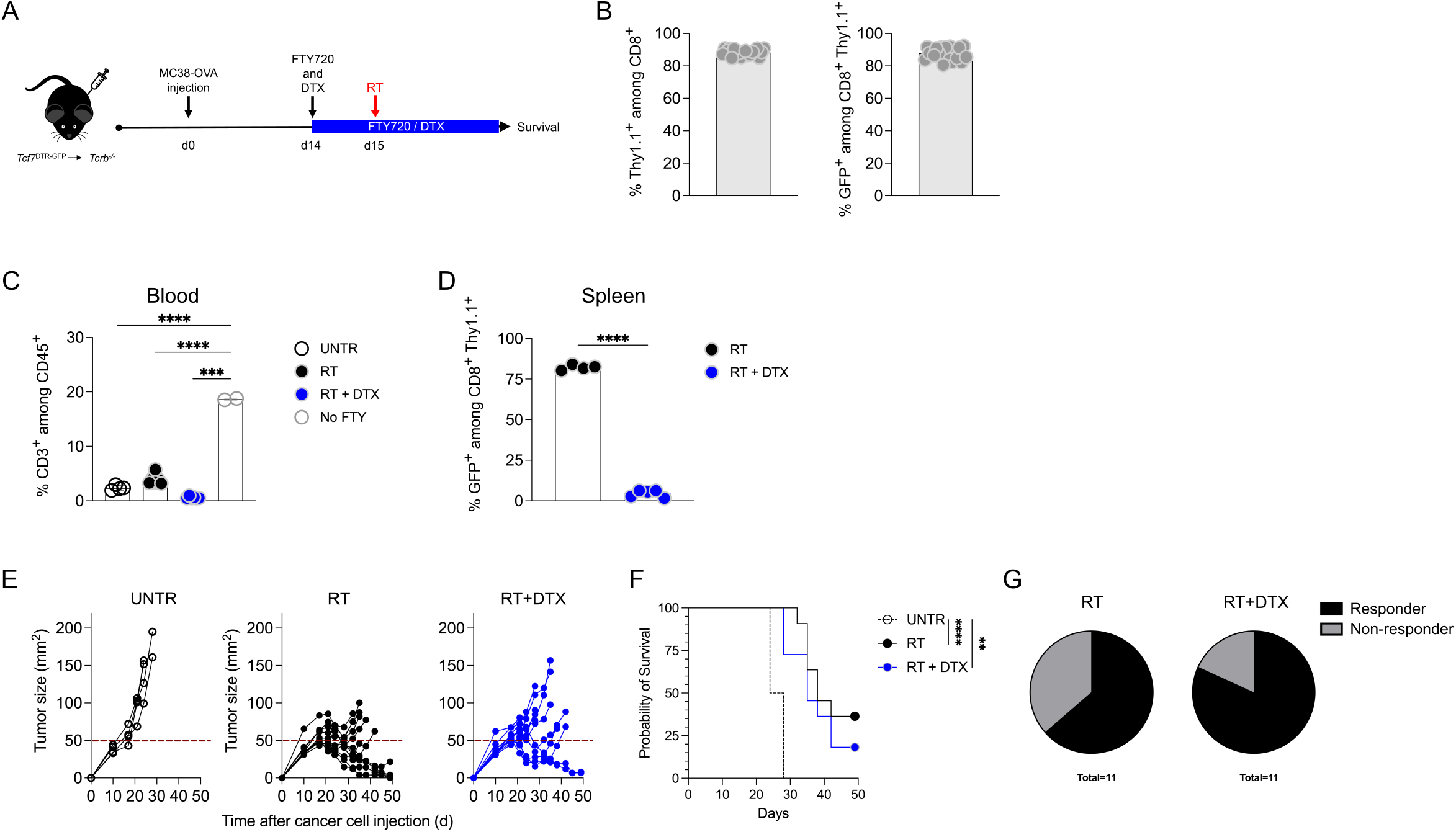
Depletion of TCF-1+ cells reduces the radiotherapy efficacy on survival. (A) Experimental setup: Tcf7DTR-GFP→Tcrb-/- mice (UNTR n=4, the rest n=11 per condition) bone marrow transplant was performed. After 8 weeks, the mice were injected subcutaneously with 8x105 MC38-OVA-GFP cells in matrigel:PBS (1:2). Fifteen days after tumor injection, some groups received radiotherapy (RT; 20 Gy) or not (UNTR). Starting 1 day before radiotherapy, all groups received 20 µg FTY720 per os every second day. Some groups received 250 ng diphtheria toxin (DTX) intraperitoneal injection twice per week, the rest received PBS until the endpoint. (B) Bone marrow transplantation success was evaluated by measuring Thy1.1+ (left) and GFP+ (right) cells among CD8+ cells in the blood one day before tumor injection. (C) FTY720 validation was performed by measuring the percentage of CD3+ cells among CD45+ cells in the blood one day after radiotherapy. (D) Depletion of TCF-1+ (GFP+) cells was validated by measuring the percentage of GFP+ cells among CD8+ cells in the spleen upon sacrifice. (E) Tumor size (length x breadth) was measured by caliper. Each symbol represents one mouse. The bar indicates the mean ± SD. Groups were statistically compared using one-way ANOVA with Tukey’s multiple comparison correction. (F) Kaplan-Meier survival curves. Groups were statistically compared using the Mantel-Cox test. (G) Pie chart showing radiotherapy responders (tumor size below 50 mm2 at day 50) and non-responders. **p<0.01, ***p<0.001, ****p<0.0001.

**Figure S14.**
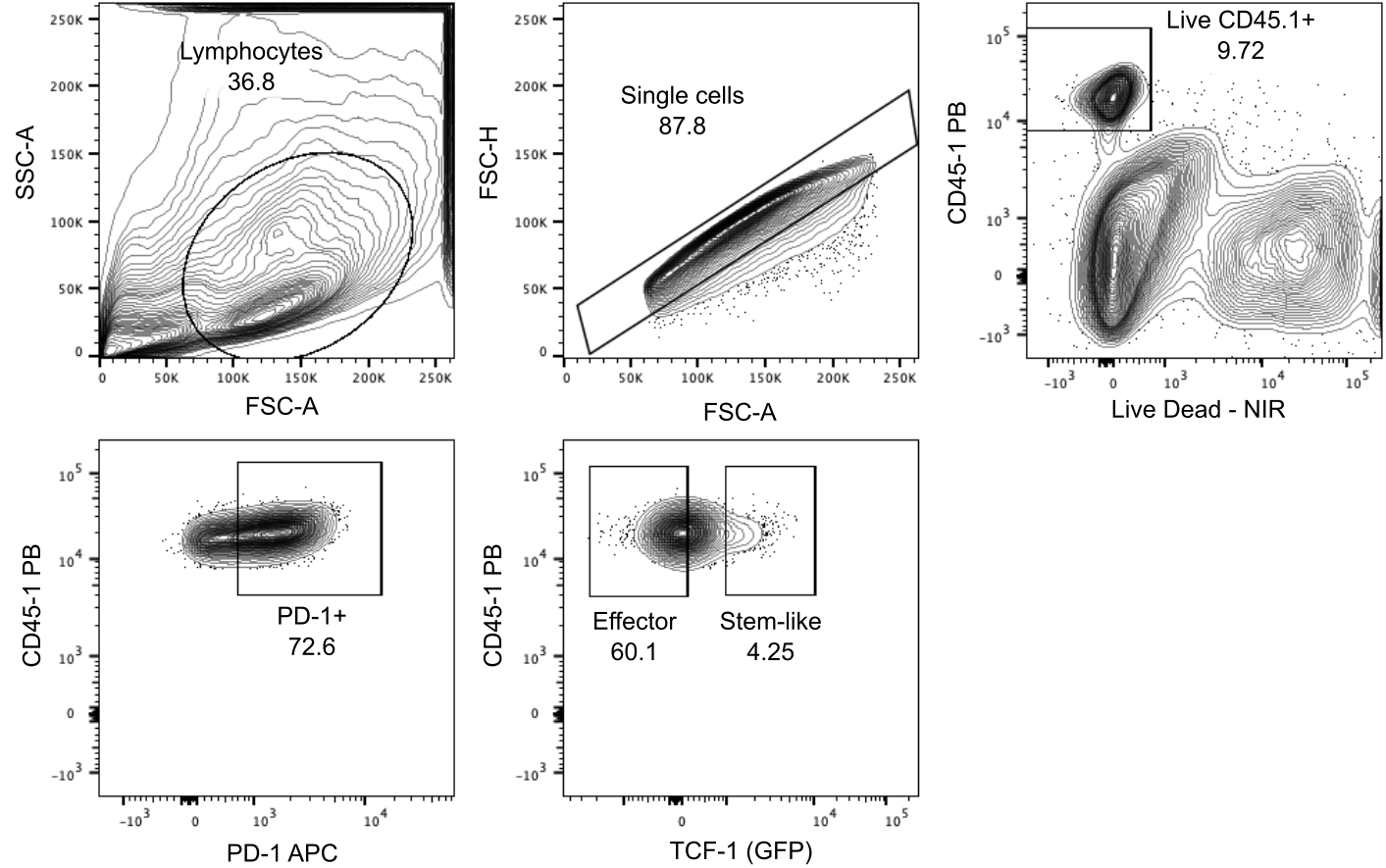
Stem-like CD8+ T-cells sorting strategy from tumor.

**Figure S15.**
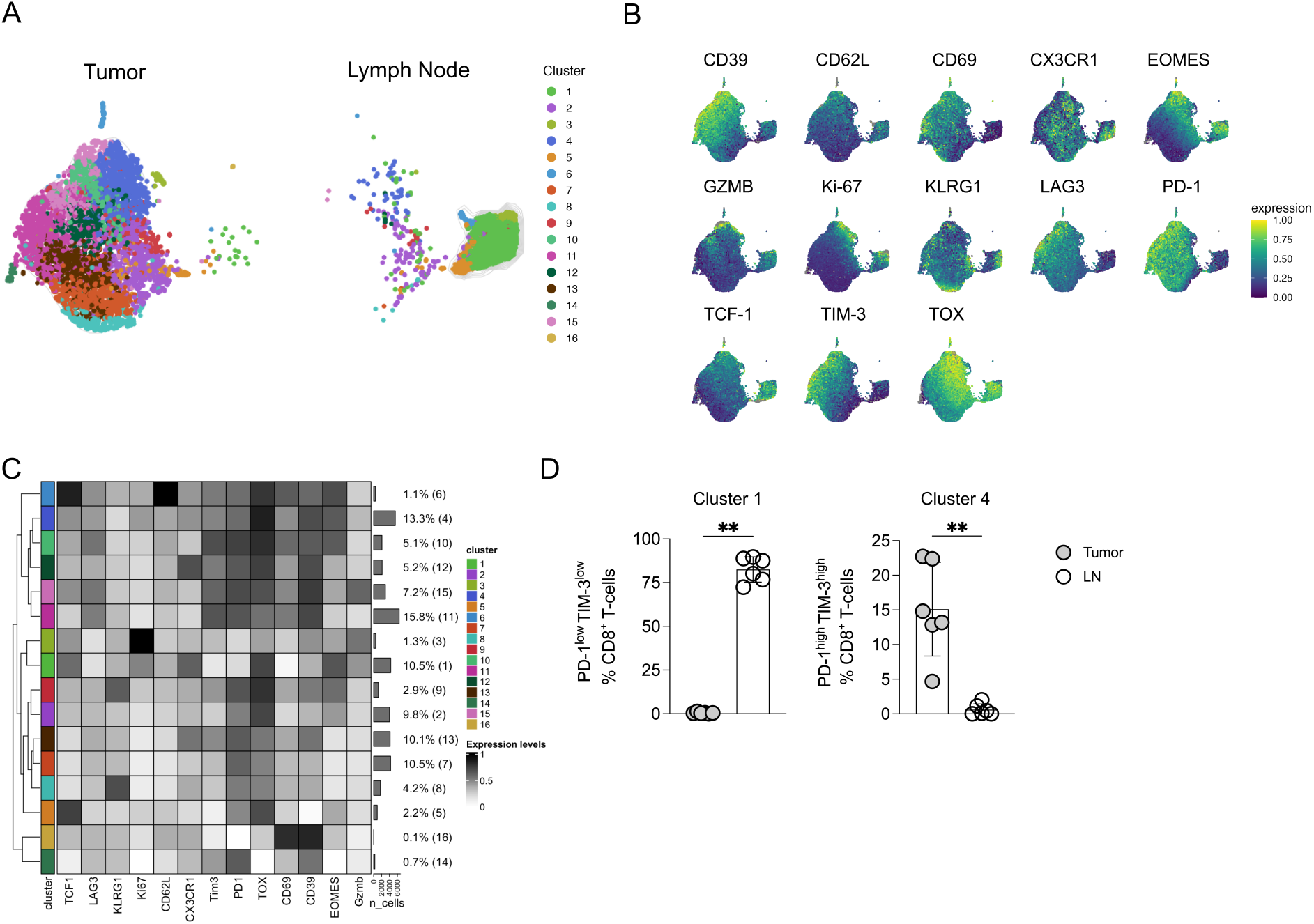
Adoptively transferred stem-like CD8+ T-cells preserve a less differentiated phenotype in the lymph node. (A) UMAP representation of CD8+ T-cell (derived from stem-like CD8+ T-cells) clusters in tumor (left) and in tumor-draining lymph node (right). (B) UMAP plots showing the expression of key markers in CD8+ T-cells. (C) The heatmap illustrates the relative expression of key markers across different clusters. (D) Percentage of UMAP clusters 1 and 2. Each symbol represents one mouse. The bar indicates the mean ± SD. Groups were statistically compared using the Mann-Whitney U test. **p<0.01.

**Figure S16.**
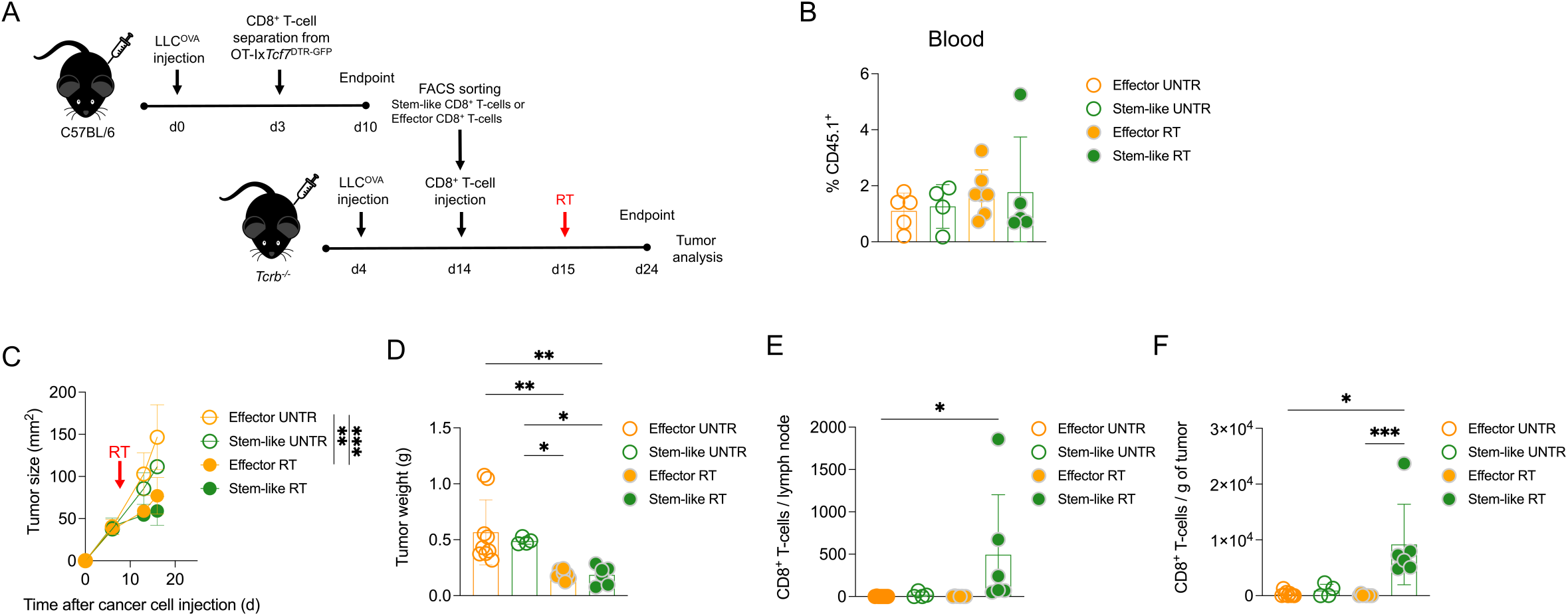
Radiotherapy is necessary for the proliferative response of stem-like CD8+ T-cells. (A) Experimental design: Stem-like CD8+ T-cells differentiate and proliferate after radiotherapy. (A) Experimental setup: C57BL/6 mice (n=15) were injected subcutaneously with 3x105 LLC-OVA cells in matrigel:PBS (1:2). Three days later, all mice received an intravenous injection of magnetically enriched OT-I x Tcf7DTR- GFP (CD45.1) CD8+ T-cells (105 cells per mouse). Seven days later, tumors were processed, and stem-like (TCF-1+PD-1+) and effector (TCF-1−PD-1+) CD8+ T-cells were sorted using flow cytometry. These cells were then intravenously injected into LLC-OVA-bearing Tcrb-/- mice (1000 cells per mouse). One day later, some mice (Effector UNTR n=9, Effector RT n=8, Stem-like UNTR n=4, Stem-like RT n=6,) received radiotherapy (RT; 20 Gy) or not (UNTR). Tumors and tumor-draining lymph nodes were analyzed nine days after radiotherapy. (B) Presence of adoptively transferred cells was evaluated by measuring the percentage of CD45.1+ cells among CD45+ cells in the blood six days after radiotherapy. (C) Tumor size (length x breadth) was measured by caliper. (D) Tumor weight (E-F) Number of CD8+ T-cells per lymph node (E) and per gram of tumor (F). Each symbol represents one mouse. The bar indicates the mean ± SD. Groups were statistically compared using one-way ANOVA with Tukey’s multiple comparison correction. *p<0.05, **p<0.01, *** p<0.001.

